# Targeted degradation of PRC1 components, BMI1 and RING1B, via a novel protein complex degrader strategy

**DOI:** 10.1101/2022.11.19.517138

**Authors:** Kwang-Su Park, Lihuai Qin, Md Kabir, Kaixiu Luo, Brandon Dale, Yue Zhong, Arum Kim, Gang Greg Wang, H. Ümit Kaniskan, Jian Jin

**Author notes:** These authors contributed equally to this work (Kwang-Su Park, Lihuai Qin, Md Kabir). **Corresponding Author** (J.J.).

## Abstract

Polycomb repressive complex 1 (PRC1) is an essential epigenetic regulator that mainly controls histone H2A Lys119 mono-ubiquitination (H2AK119ub). BMI1 and RING1B are PRC1 core components and play critical roles in the development of various cancers. However, therapeutic agents targeting PRC1 are very limited, and small-molecule inhibitors of PRC1 displayed limited effectiveness in killing cancer cells. In this study, MS147, the first degrader of PRC1 core components, BMI1 and RING1B, was discovered via a novel protein complex degradation strategy that utilizes the target protein’s interacting partner protein (EED) to degrade BMI1 and RING1B. MS147, which comprises an EED small-molecule binder linked to a ligand of the E3 ligase VHL, degrades BMI1 and RING1B in an EED-, VHL-, ubiquitination- and time-dependent manner. MS147 is selective and preferentially degrades BMI1 and RING1B over PRC2 core components: EED, EZH2 and SUZ12. Consequently, MS147 effectively reduces H2AK119ub, but not H3K27me3, which is catalyzed by PRC2. Furthermore, MS147, but not the parent EED binder or known PRC2 degraders, effectively inhibits the proliferation of cancer cell lines that are insensitive to EZH2 knockout or PRC2 degraders. Overall, this study provides a novel degrader targeting BMI1 and RING1B, which is a useful chemical tool to further investigate the roles of PRC1 in cancer, and a novel protein complex degradation strategy, which could potentially expand the degradable human proteome.

## 1. Introduction

Polycomb group (PcG) proteins, which include polycomb repressive complex 1 (PRC1) and 2 (PRC2), have been well studied as key histone modifiers to regulate gene transcription.^[1-3]^ PRC1 is a multi-subunit protein complex, containing RING1A or RING1B with one of the six PCGF1-PCGF6 paralogs, which catalyzes mono-ubiquitination of histone H2A lysine 119 (H2AK119ub).^[4]^ As core components of the main heterodimeric complex of canonical PRC1, BMI1 (PCGF4) and RING1B (RNF2) are sufficient for catalyzing H2AK119ub.^[5]^ On the other hand, PRC2, which consists of EZH2, SUZ12 and EED as core components, is the methyltransferase responsible for trimethylation of histone H3 lysine 27 (H3K27me3).^[6]^

Recent studies support that both enzymatic and non-enzymatic functions of PRC1 core components, especially BMI1 and RING1B, are critically involved in the development and progression of various tumor types.^[7]^ In particular, BMI1 is overexpressed in many different types of cancer including breast,^[8]^ lung^[9]^ and blood cancer.^[10]^ RING1B is also highly involved in breast cancer malignancy^[11]^ and leukemia progression.^[12]^ Therefore, targeting PRC1 core components could provide a potential therapeutic approach for treating cancers with alterations in PRC1 components.

To date, only two PRC1 small-molecule inhibitors, PTC209^[13]^ and RB-3,^[14]^ have been reported. However, direct interaction between PTC209 and any PRC1 components has not been established to support that PTC209 is a direct inhibitor of PRC1. RB-3 displayed very limited effectiveness in killing various types of cancer cells.^[14]^ Thus, a novel therapeutic approach that targets PRC1 more effectively is desirable.

Proteolysis targeting chimeras (PROTACs) are heterobifunctional small molecules that induce targeted protein degradation through hijacking the ubiquitin-proteasome system (UPS).^[15-23]^ PROTACs, as a new class of therapeutic modalities, can eliminate both catalytic and non-catalytic functions of the target enzymes or enzyme complexes. For example, we recently developed an EZH2 PROTAC degrader, which effectively targeted both canonical and non-canonical functions of EZH2, resulting in far superior tumor suppressive effects compared to catalytic inhibitors of EZH2.^[24]^

Several protein complex degraders mediated through the traditional PROTAC mechanism have been developed.^[25-31, 24]^ For example, previously reported EED PROTACs induced degradation of other PRC2 components in addition to EED.^[26, 25]^ It is important to note that none of these traditional protein complex PROTACs preferentially degraded partner proteins over the protein that binds the PROTAC directly. We thought, however, a new protein complex degrader strategy, which preferentially degrades interacting partner proteins over the protein which directly binds the PROTAC, is possible. Specifically, we hypothesized that PRC1 core components, BMI1 and RING1B, could be preferentially degraded over EED by an EED-binding PROTAC with an appropriate linker and an E3 ligase ligand (Figure 1a). In addition to being one of the core components of PRC2, EED has been reported to interact with PRC1 core components.^[32-33]^ Using immunoprecipitation (IP), we also validated the interaction between EED and PRC1 core components, BMI1 and RING1B (Figure S1).

**Figure 1.**
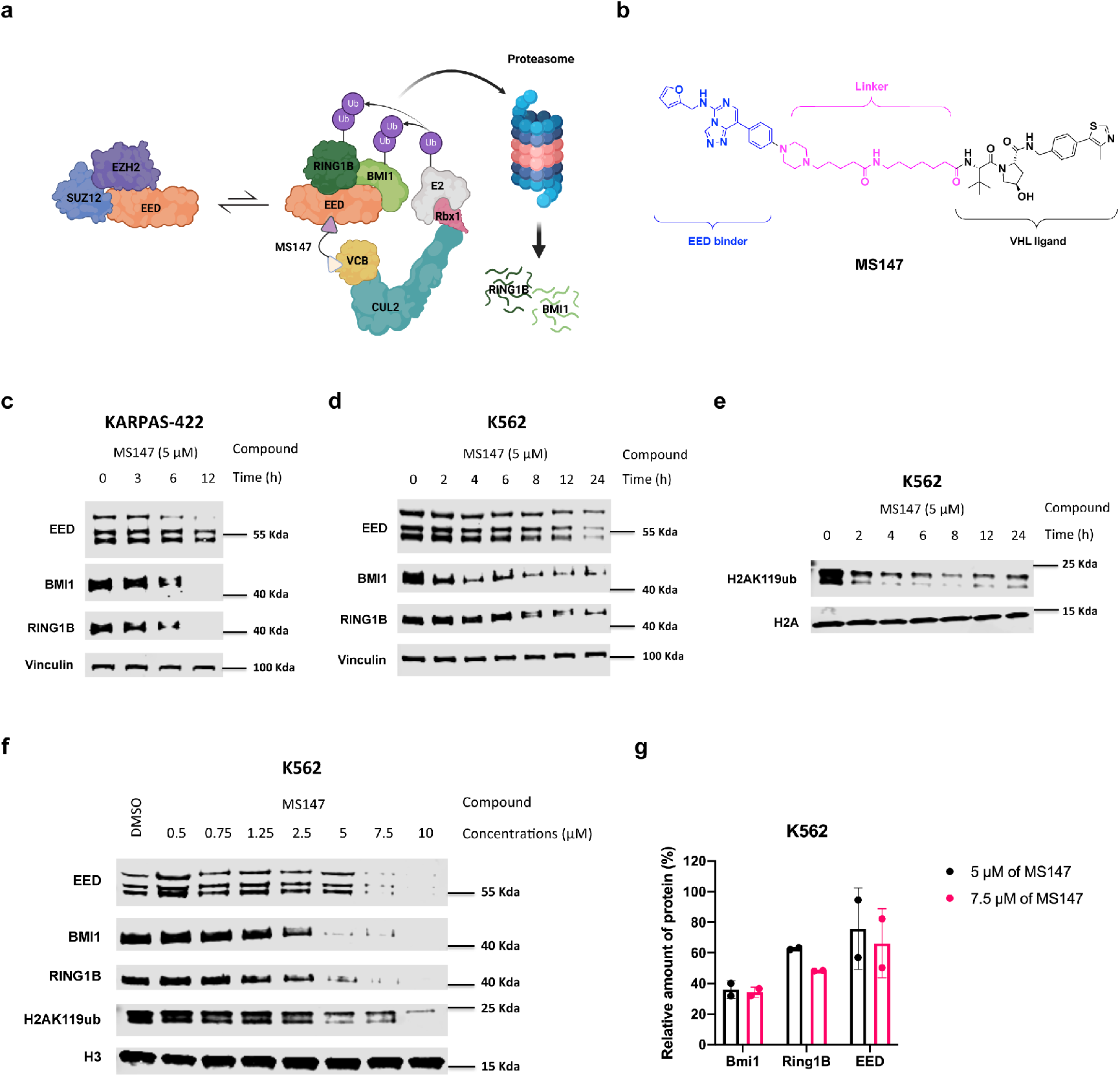
Discovery of the BMI1 and RING1B PROTAC MS147, which preferentially degrades BMI1 and RING1B over EED. (a) The schematic of an EED-binding PROTAC that preferentially degrades PRC1 components, BMI1 and RING1B, over EED, by hijacking the VHL-Elongin C-Elongin B (VCB) cullin-2 (CUL2) RING E3 ligase complex. (b) Chemical structure of MS147. (c-d) Time-dependent degradation of BMI1, RING1B and EED induced by MS147 in KARPAS-422 (c) and K562 (d) cells. KARPAS-422 and K562 cells were treated with MS147 at 5 μM for the indicated time. The protein levels of EED, BMI1 and RING1B were determined by Western blotting (WB) with vinculin as the loading control. (e) Time course of the H2AK119ub reduction induced by MS147 in K562 cells. K562 cells were treated with MS147 at 5 μM for the indicated time. The H2AK119ub protein level was determined by WB with H2A as the loading control. (f) Concentration-dependent degradation of BMI1, RING1B and EED and reduction of H2AK119ub induced by MS147 in K562 cells. K562 cells were treated with MS147 at the indicated concentrations for 24 h. The protein levels of EED, BMI1, RING1B and H2AK119ub were determined by WB with vinculin and H3 as the loading controls. (g) Quantification of the EED, BMI1 and RING1B protein levels in K562 cells treated with 5 or 7.5 μM of MS147 for 24 h. The WB results shown in panels c-f are representative of two independent experiments.

Using this new protein complex degrader strategy, we discovered the first BMI1 and RING1B degrader, MS147, which consists of an EED binder linked to a VHL E3 ligase ligand. MS147 preferentially degraded BMI1 and RING1B over EED and other PRC2 components in an EED-, VHL- and ubiquitination-dependent manner. Consequently, MS147 effectively decreased H2AK119ub, but not H3K27me3. Phenotypically, MS147, but not the parent EED binder or published EED/PRC2 degraders, effectively inhibited the proliferation in multiple cancer cell lines that are insensitive to either EZH2 knockout or EED/PRC2 degraders. Overall, this study presents a novel protein complex degradation strategy and a novel BMI1 and RING1B degrader, which could be a valuable chemical tool for the scientific community to further study PRC1.

## 2. Results

### 2.1. Discovery of MS147, which preferentially degrades PRC1 components, BMI1 and RING1B, over EED

To demonstrate the applicability of the novel protein complex degradation strategy (Figure 1a), we designed a set of putative BMI1/RING1B degraders using a highly potent and selective small-molecule binder of EED, EED226, which allosterically inhibits PRC2.^[34]^ By analyzing the co-crystal structure of EED in complex with EED226 (PDB ID: 5WUK)^[35]^, we identified a solvent-exposed region to conjugate a linker (Figure S2a). We modified the solvent-exposed methylsulfonyl moiety in EED226 to a piperazinyl moiety, which was conjugated to VHL1, a well-known ligand of the E3 ligase von Hippel-Lindau (VHL)^[15]^, via various linkers (Figure S2b). We chose to hijack the E3 ligase VHL, instead of cereblon (CRBN), mainly to avoid the potential complication caused by CRBN neosubstrate degradation. We next determined the effect of these compounds on reducing the protein levels of BMI1, RING1B, EED and H2AK119ub in K562 cells, a chronic myelogenous leukemia cell line (Figure S3). From this study, we identified compound **6** (MS147, Figure 2b) as a promising BMI1 and RING1B PROTAC, which effectively degraded BMI1, RING1B and EED and reduced H2AK119ub in K562 cells treated with the compound for 24 h (Figure S3).

**Figure 2.**
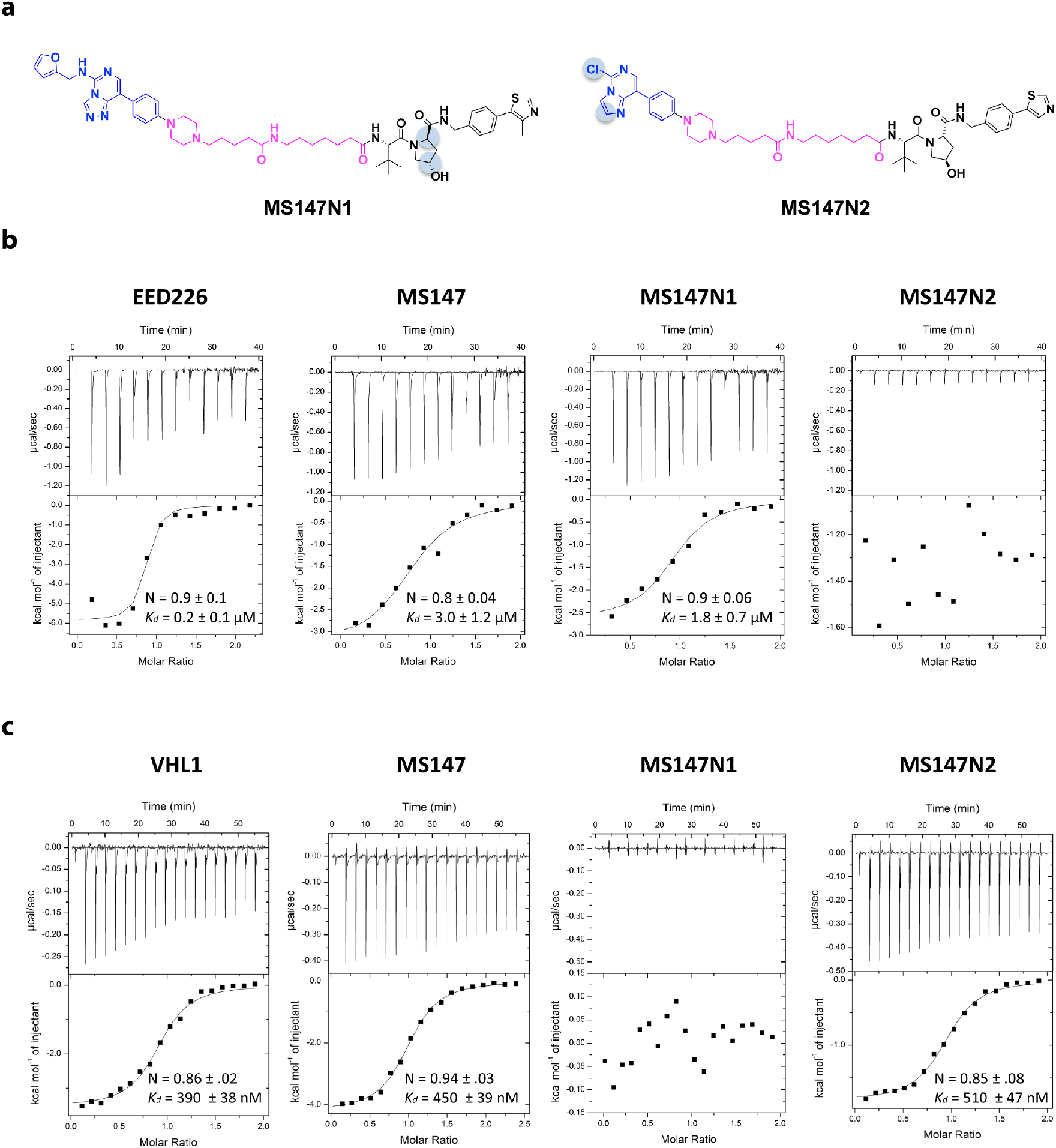
MS147 binds both EED and VHL while the negative control MS147N1 binds EED but not VHL, and MS147N2 binds VHL but not EED. (a) Chemical structures of the negative controls, MS147N1 and MS147N2. (b) Binding affinities of MS147, MS147N1 and MS147N2 to EED were determined using ITC. EED226 was used as a positive control. (c) Binding affinities of MS147, MS147N1 and MS147N2 to VHL were determined using ITC. VHL1 was used as a positive control. The calculated values in panels b and c represent the means ± SD from two independent experiments.

We next characterized the degradation profile of MS147 in K562 and KARPAS-422 (B-cell lymphoma) cell lines. These cell lines were chosen mainly due to the association of PRC1 with the hematopoiesis process in blood cancers^[36]^. In time course studies, we found that MS147 degraded BMI1 and RING1B effectively as early as 6 hours, and almost completely at 12 hours, while it degraded EED partially at 12 hours in KARPAS-422 cells (Figure 1c). Similarly, in K562 cells, MS147 induced degradation of BMI1 as early as 4 hours and RING1B as early as 8 hours, while it induced modest EED degradation even at 12 hours (Figure 1d). These results indicate that MS147 preferentially degrades BMI1 and RING1B over EED. Furthermore, MS147 decreased the H2AK119ub level in a time-dependent manner in K562 cells (Figure 1e). MS147 also concentration-dependently degraded BMI1 and RING1B and reduced H2AK119ub in K562 cells (Figure 1f). While MS147 also degraded EED at high concentrations (7.5 and 10 μM) after 24 hours of treatment, MS147 at 5 and 7.5 μM preferentially degraded BMI1 and RING1B over EED (Figure 1g). Overall, our design, structure-activity relationship, and initial characterization studies resulted in the discovery of MS147, which preferentially degrades PRC1 components, BMI1 and RING1B, over EED.

### 2.2. MS147 degrades BMI1 and RING1B in an EED-, VHL- and ubiquitination-dependent manner

To confirm the mechanism of degradation of MS147, we first developed 2 close analogs of MS147, MS147N1 and MS147N2, as negative controls of MS147 (Figure 2a). MS147N1, which contains the same EED binder and linker but a diastereomer of the VHL ligand,^[30]^ was designed to bind EED but not VHL. MS147N2, which contains the same VHL ligand and linker but a modified EED binding moiety,^[37]^ was designed to maintain the binding to VHL but not EED. Using isothermal titration calorimetry (ITC), we confirmed that MS147 (*K*_d_ = 3.0 ± 1.2 μM) and MS147N1 (*K*_d_ = 1.8 ± 0.7 μM) bound EED with similar binding affinities, while MS147N2 did not bind EED (Figure 2b). Although binding affinities of MS147 and MS147N1 to EED are lower than that of EED226, this level of binding affinities is sufficient for PROTACs to induce effective targeted protein degradation.^[18]^ We also confirmed that MS147 (*K*_d_ = 450 ± 39 nM) and MS147N2 (*K*_d_ = 510 ± 47 nM) showed similar binding affinities for VHL, while MS147N1 did not bind VHL (Figure 2c). These ITC results indicate that MS147 directly binds EED and VHL, while the negative control MS147N1 binds EED but not VHL, and MS147N2 binds VHL but not EED.

To confirm that the degradation of BMI1 and RING1B induced by MS147 is mediated through EED and VHL, we conducted a number of mechanism of action (MOA) studies in K562 cells. We first confirmed that MS147N1, which binds EED but not VHL, and MS147N2, which binds VHL but not EED, did not degrade BMI1, RING1B and EED, while MS147 did (Figure 3a). We next generated K562 cells with EED knockdown (KD) using shEED and further assessed the dependency of the BMI1 and RING1B degradation induced by MS147 on EED. As expected, the ability of MS147 in degrading BMI1 and RING1B was completely abolished upon EED depletion (Figure 3b). Notably, EED depletion alone (without MS147 treatment) did not lead to degradation of BMI1 and RING1B (Figure 3b), suggesting that the BMI1 and RING1B degradation effect of MS147 is not a consequence of EED degradation. Together, these results indicate that the BMI1 and RING1B degradation induced by MS147 is dependent on EED.

**Figure 3.**
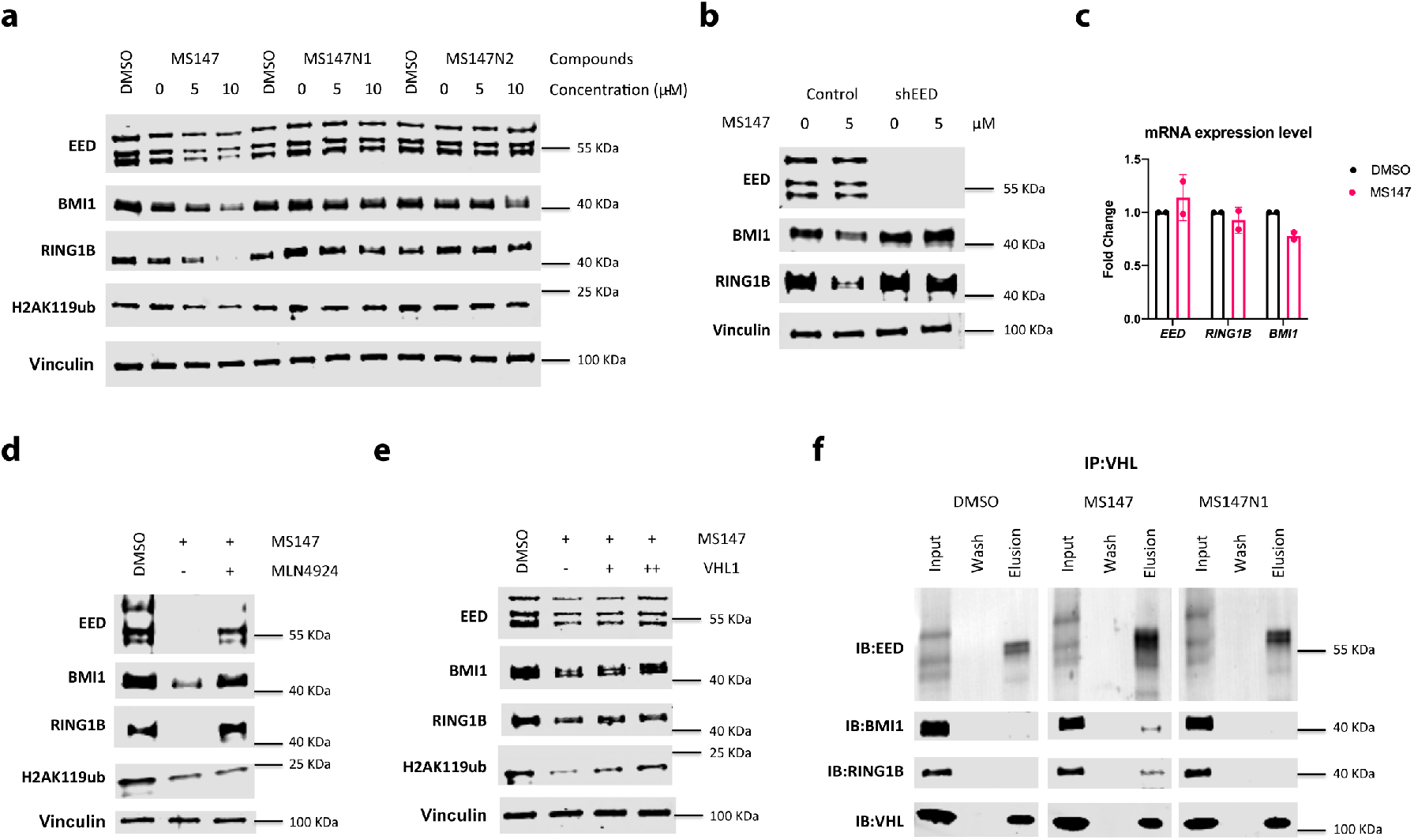
MS147 degrades BMI1 and RING1B in an EED-, VHL- and ubiquitination-dependent manner without altering mRNA expression levels. (a) The effect of MS147, MS147N1 and MS147N2 on reducing the protein levels of EED, BMI1, RING1B and H2AK119ub. K562 cells were treated with DMSO or the indicated compound at the indicated concentrations for 24 h. (b) The effect of EED knockdown (KD) using shEED on rescuing MS147-induced degradation of BMI1 and RING1B. K562 cells were transfected by lentivirus containing shEED or an empty vector for 24 h. The transfected cells were then treated with DMSO or MS147 (5 μM) for 24 h. (c) The effect of MS147 on the mRNA levels of *EED, BMI1* and *RING1B*, determined by RT-qPCR. KARPAS-422 cells were treated with DMSO or MS147 (5 μM) for 24 h. The mRNA levels were normalized to DMSO control. The data shown represent the means ± SD from two independent experiments. (d-e) The effect of pretreatment with the neddylation inhibitor MLN4924 (d) or the VHL ligand VHL1 (e) on rescuing the degradation of BMI1 and RING1B induced by MS147. K562 cells were pretreated with MLN4924 (0.5 μM) or VHL1 (+: 0.5 μM; ++: 1 μM) for 1 h, followed by treatment with MS147 (5 μM) for 24 h. (f) Co-elution of EED-BMI1-RING1B with VHL by *in vitro* pulldown using a VHL antibody in the presence of MS147 or MS147N1. K562 cell lysates were treated with DMSO, MS147 (40 μM) or MS147N1 (40 μM) for 3 h. The protein levels of EED, BMI1, RING1B and/or H2AK119ub in panels a, b, and d-f were determined by WB with vinculin as the loading control for a, b, d and e. The WB results shown in panels a, b, and d-f are representative of two independent experiments.

To further validate that the BMI1 and RING1B degradation induced by MS147 occurs through recruiting the VHL-mediated ubiquitination system, we next performed RT-qPCR, additional rescue and *in vitro* pulldown experiments. First, we evaluated the effect of MS147 on the mRNA expression levels of *BMI1, RING1B* and *EED* in KARPAS-422 cells, in which MS147 induced robust degradation of BMI1 and RING1B (Figure 1c). As shown in Figure 3c, MS147 treatment did not significantly change the mRNA expression levels of *BMI1, RING1B* and *EED*, indicating that the observed BMI1 and RING1B degradation induced by MS147 is not due to changes in transcription. Next, we performed additional rescue experiments by pretreating K562 cells with a neddylation inhibitor or a VHL binder. Pretreatment with MLN4924,^[38]^ an inhibitor of the NEDD8-activating enzyme (NAE), which is responsible for neddylation of cullin-RING E3 ubiquitin ligases, rescued the BMI1 and RING1B degradation effect induced by MS147 (Figure 3d). By competing with MS147 for VHL binding, VHL1, the VHL binder used in MS147, also rescued MS147’s BMI1 and RING1B degradation effect (Figure 3e). Interestingly, the level of H2AK119ub was also recovered by pretreatment with VHL1, but not MLN4924. This result is likely due to the ubiquitination inhibition effect of MLN4924, which prevents the re-ubiquitination of H2A. Lastly, we conducted *in vitro* pulldown experiments to assess the formation of the BMI1-RING1B-EED-MS147-VHL complex in K562 cells. We were pleased to find that BMI1, RING1B and EED co-eluted with VHL in the presence of MS147, but not MS147N1 (Figure 3f), indicating that MS147 recruits VHL to EED and its interacting partners BMI1 and RING1B.

Collectively, the results of these MOA studies indicate that the degradation of BMI1 and RING1B induced by MS147 is dependent on EED, VHL and ubiquitination. The induced degradation occurs through the formation of the BMI1-RING1B-EED-MS147-VHL complex without altering mRNA expression levels of *BMI1* and *RING1B*. These results support MS147’s mechanism of degradation outlined in Figure 1a.

### 2.3. MS147 is selective for PRC1 over PRC2 and for EED over other epigenetic proteins

We next assessed selectivity of MS147 for PRC1 over PRC2 and for EED over a panel of 20 methyltransferases and 12 epigenetic reader proteins. First, since EED is a core component of PRC2, we evaluated the degradation effect of MS147 on other PRC2 core components, EZH2 and SUZ12, in K562 cells. EED226 and the previously published EED/PRC2 degrader, PROTAC 2^[25]^, were used as controls. It was reported previously that PROTAC 2 effectively degraded EED first and other PRC2 components, EZH2 and SUZ12, subsequently.^[25]^ The degradation of EZH2 and SUZ12 by PROTAC 2 is likely a consequence of EED degradation, due to the destabilization of PRC2 after losing one of its core components (illustrated in Figure S4). To confirm this, we performed KD experiments using siEED. Indeed, depletion of EED substantially reduced EZH2 and SUZ12 protein levels (Figure S5), suggesting that PROTAC 2-mediated EZH2 and SUZ12 degradation is a consequence of the PRC2 destabilization due to EED degradation.

Importantly, MS147 selectively degraded BMI1 and RING1B over PRC2 core components, EZH2 and SUZ12, in addition to EED in K562 cells (Figures 1 and 4a-c), in contrast to PROTAC 2, which robustly degraded PRC2 components, EED, EZH2 and SUZ12, but not PRC1 components, BMI1 and RING1B (Figure 4a). As expected, EED226 did not degrade any of PRC1/PRC2 components (Figure 4a), indicating that EED binding itself has no impact on the degradation effect of MS147 and PROTAC 2. To further evaluate the degradation effect of MS147 on EZH2 and SUZ12, we monitored EZH2 and SUZ12 protein levels after treatment of MS147 in K562 cells at different concentrations and time points (Figure 4b-c). While MS147 induced modest degradation of EZH2 and SUZ12 at high concentrations such as 10 μM (Figure 4b), it displayed little effect on degrading EZH2 and SUZ12 in the time course study (Figure 4c). MS147’s modest degradation of EZH2 and SUZ12 could be due to its degradation effect on EED, since MS147 degraded EED faster than EZH2 and SUZ12 (Figures 1c-d and 4c) and EED KD resulted in drastic reduction in the protein levels of EZH2 and SUZ12 (Figure S5). Overall, MS147 selectively degrades BMI1 and RING1B over EED, EZH2 and SUZ12 (Figures 1 and 4a-c). Another evidence supporting MS147 having a very modest effect on PRC2 is that while EED226 and PROTAC 2 effectively reduced H3K27me3, the product of PRC2’s methyltransferase activity, MS147 minimally decreased H3K27me3 (Figure 4a-c). Instead, MS147 significantly reduced H2AK119ub (Figure 1e-f), which is mediated by PRC1. We also evaluated the degradation effect of MS147 on RING1A, which is highly homologous to RING1B. Not surprisingly, MS147 also induced degradation of RING1A in K562 cells (Figure S6). Since it has been reported that RING1B is predominantly responsible for mono-ubiquitination of H2A instead of RING1A^[39]^, we mainly focused on RING1B in this study. Collectively, our results support that MS147 selectively degrades PRC1 components, BMI1 and RING1B, over PRC2 components, EED, EZH2 and SUZ12, and selectively reduces H2AK119ub (mediated by PRC1) over H3K27me3 (mediated by PRC2).

**Figure 4.**
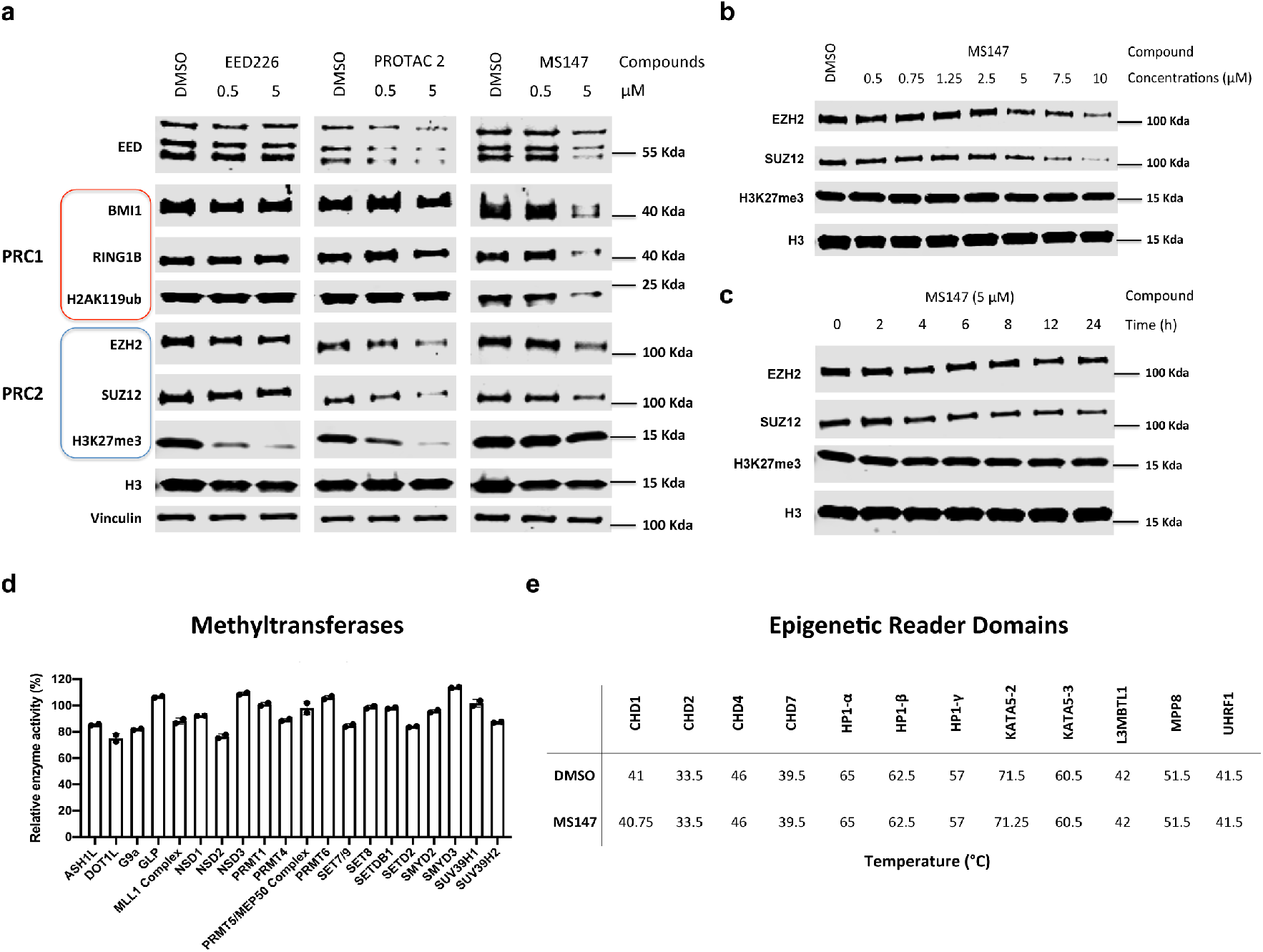
MS147 selectively degrades BMI1 and RING1B over PRC2 core components and is selective for EED over 20 methyltransferases and 12 epigenetic reader domains. (a) The effect of MS147, PROTAC 2 and EED226 on reducing the protein levels of EED, BMI1, RING1B, H2AK119ub, EZH2, SUZ12 and H3K27me3. K562 cells were treated with DMSO or the indicated compound at the indicated concentrations for 24 h. (b-c) The effect of MS147 on reducing the EZH2, SUZ12 and H3K27me3 protein levels in K562 cells treated with MS147 at a range of concentrations for 24 h (b) or 5 μM of MS147 for indicated times (c). The protein levels of EED, BMI1, RING1B, H2AK119ub, EZH2, SUZ12 and/or H3K27me3 in panels a-c were determined by WB with vinculin and/or H3 as the loading controls. The WB results shown in panels a-c are representative of two independent experiments. (d-e) The effect of MS147 on inhibiting 20 methyltransferases in biochemical assays (d) and binding 12 epigenetic reader domains in *in vitro* thermal shift assays (e). MS147 was tested in these selectivity assays at 10 μM. Data shown are the means ± SD from duplicate experiments.

**Figure 5.**
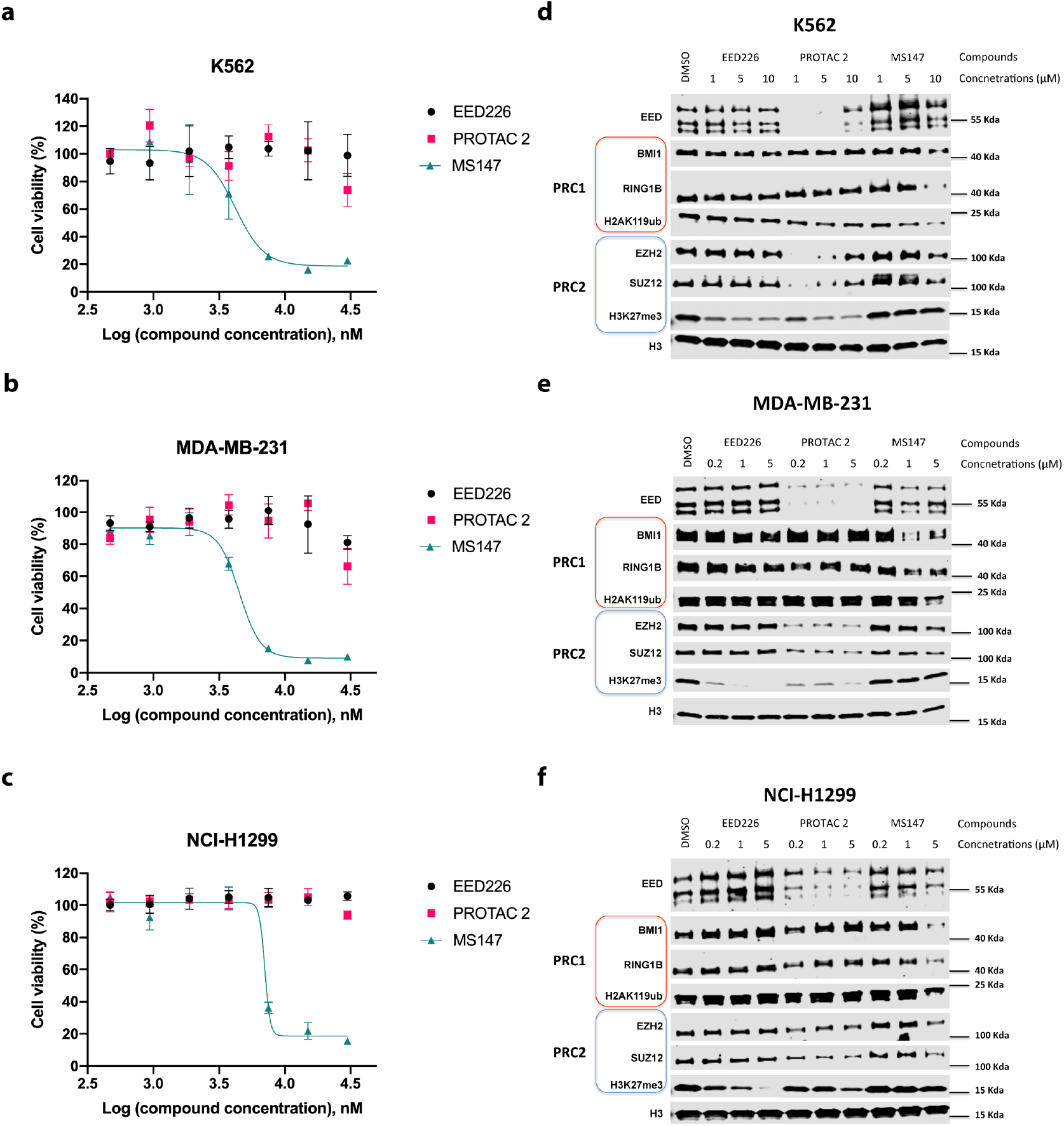
MS147, but not the parent EED binder, EED226, or EED/PRC2 degrader, PROTAC 2, effectively suppresses the proliferation in K562, MDA-MB-231 and NCI-H1299 cells. (a-c) The effect of MS147, EED226 and PROTAC 2 on inhibiting the growth in K562 (a), MDA-MB-231 (b) and NCI-H1299 (c) cells. The tested cell lines were treated with serial dilution of EED226, PROTAC 2 or MS147 for 5 days. Cell viability was determined using the CCK-8 assay. The data shown represent the means ± SD from three independent experiments. (d-f) The effect of MS147, EED226 and PROTAC 2 on degrading BMI1, RING1B, EED, EZH2 and SUZ12 and reducing the levels of H2AK119ub and H3K27me3 in K562 (d), MDA-MB-231 (e) and NCI-H1299 (f) cells. The above three cell lines were treated with EED226, PROTAC2 or MS147 at the indicated concentrations for 24 h (d) or 48 h (e, f). The protein levels of EED, BMI1, RING1B, H2AK119ub, EZH2, SUZ12 and H3K27me3 were determined using WB with H3 as the loading control. The WB results shown are representative of two independent experiments.

We next evaluated selectivity of MS147 against a panel of 20 methyltransferases and 12 epigenetic reader domains (Figure 4d-e). The enzymatic inhibition effect of MS147 at 10 μM against 20 methyltransferases was determined using radioactivity-based biochemical assays. As expected, MS147 did not show significant inhibition effect (<30% at 10 μM) against any of the 20 methyltransferases (Figure 4d). Binding of MS147 to 12 epigenetic reader domains was also assessed using *in vitro* thermal shift assays. MS147 at 10 μM did not display notable melting temperature (Tm) changes against these 12 reader domains (Figure 4e). Together, these results support that MS147 is selective for EED over other epigenetic modulating proteins.

### 2.4. MS147, but not the parent EED binder or EED/PRC2 degrader, effectively inhibits the proliferation in multiple cancer cell lines

We next investigated the anti-proliferative activity of MS147 in 3 different cancer cell lines: K562 (leukemia), MDA-MB-231 (triple-negative breast cancer), NCI-H1299 (non-small cell lung cancer); and its toxicity on the PNT2 normal prostate cell line. We selected these cell lines because it has been shown that the growth of K562 and MDA-MB-231 cells is insensitive to EZH2 knockout via CRISPR^[40-41]^, and the growth of NCI-H1299 and PNT2 cells is insensitive to EED/PRC2 or EZH2 degraders^[41, 25]^. We first confirmed that K562, MDA-MB-231 and NCI-H1299 cells were indeed insensitive to the parent EED binder, EED226, and EED/PRC2 degrader, PROTAC 2 (Figure 6a-c). On the other hand, MS147 effectively suppressed the proliferation in K562 (GI_50_ = 3.8 ± 1.0 μM), MDA-MB-231 (GI_50_ = 4.5 ± 0.2 μM) and NCI-H1299 (GI_50_ = 6.8 ± 0.5 μM) cells in a concentration-dependent manner (Figure 6a-c).

We next determined the effect of MS147, EED226 and PROTAC 2 on degrading PRC1 and PRC2 components in K562, MDA-MB-231 and NCI-H1299 cells. As shown in Figure 6d-f, MS147 preferentially degraded PRC1 core components, BMI1 and RING1B, over PRC2 core components: EED, EZH2 and SUZ12, in all three cell lines. Furthermore, MS147 effectively reduced the PRC1-mediated H2AK119ub mark without changing the PRC2-mediated H3K27me3 mark in all three cell lines. On the other hand, PROTAC 2 robustly degraded PRC2 core components, EED, EZH2 and SUZ12, but not PRC1 core components, BMI1 and RING1B, in these three cell lines. Consequently, PROTAC 2 effectively decreased H3K27me3, but not H2AK119ub. As expected, EED226 did not degrade any PRC1/PRC2 components, and it effectively reduced H3K27me3, but not H2AK119ub, in these cells. Collectively, these results suggest that the anti-proliferative effect of MS147 in K562, MDA-MB-231 and NCI-H1299 cells is mainly due to the effect of MS147 on degrading BMI1 and RING1B, but not its modest effect on degrading PRC2 core components.

Finally, we assessed the toxicity of MS147 using a normal prostate cell line, PNT2, and found that MS147 did not display any cell growth inhibition effect even at high concentrations such as 30 μM (Figure S7). The parent EED binder, EED226, and EED/PRC2 degrader, PROTAC 2, were also not toxic to PNT2 cells.

Taken together, these results suggest that BMI1/RING1B degraders, such as MS147, but not the parent EED binder or EED/PRC2 degraders, can effectively suppress the proliferation of cancer cell lines that are insensitive to EZH2 knockout or EED/PRC2 degraders. Our results also support that MS147 is not toxic in normal cells.

## 3. Discussion

In this study, we discovered the first BMI1 and RING1B degrader, MS147, using a novel protein complex degrader strategy, which utilizes a small-molecule ligand that binds EED, an interacting partner protein of BMI1 and RING1B, to recruit BMI1 and RING1B to the E3 ligase VHL for ubiquitination and degradation by the UPS. We hypothesized that PRC1 core components BMI1 and RING1B could be preferentially degraded over EED and other PRC2 core components, EZH2 and SUZ12, by an EED-binding PROTAC with an appropriate linker and E3 ligase ligand. By conducting a structure-activity relationship study that focused on exploring various linkers, we identified MS147, which preferentially degraded PRC1 components, BMI1 and RING1B, over PRC2 components: EED, EZH2 and SUZ12, in multiple cancer cell lines. Consequently, MS147 effectively reduced the H2AK119ub mark, which is catalyzed by PRC1, without altering the H3K27me3 mark, which is catalyzed by PRC2. MS147 was also selective for EED over a panel of >30 methyltransferases and epigenetic reader proteins. Phenotypically, MS147 effectively suppressed the proliferation in multiple cancer cell lines that are insensitive to EZH2 genetic deletion or EED/PRC2 degraders, without toxic effect on normal cells.

By conducting a number of MOA studies, we confirmed that the MS147-mediated BMI1 and RING1B degradation occurs in an EED-, VHL- and ubiquitination-dependent manner, without altering mRNA expression levels of *BMI1* and *RING1B*. We show that MS147 is capable of inducing the formation of the BMI1-RING1B-EED-MS147-VHL complex, thus bringing BMI1 and RING1B in close proximity to VHL for ubiquitination and subsequent degradation. We also developed two negative controls, MS147N1 and MS147N2, which have very high structural similarity to MS147. MS147N1, which binds EED but not VHL, and MS147N2, which binds VHL but not EED, did not degrade BMI1 and RING1B nor reduce the H2AK119ub level, further supporting the EED- and VHL-dependent MOA for the BMI1 and RING1B degradation induced by MS147.

While our BMI1 and RING1B degrader, MS147, and the EED/PRC2 degrader, PROTAC 2, share similar VHL ligand and EED binder, MS147 and PROTAC 2 differ in their linkers. This linker difference is likely the main contributor to the difference in their degradation profiles. PROTAC 2 is a traditional protein complex degrader, which degraded EED – the protein it binds directly – first, and EZH2 and SUZ12 subsequently. The degradation of EZH2 and SUZ12 induced by PROTAC 2 is likely a consequence of the destabilization of PRC2 due to EED degradation, which is supported by the fact that genetic depletion of EED resulted in near complete loss of EZH2 and SUZ12. Notably, PROTAC 2 did not degrade BMI1 and RING1B. On the other hand, MS147 represents a new type of protein complex degraders. MS147 degraded BMI1 and RING1B faster and more effectively than EED, the protein it binds directly, and other PRC2 components (EZH2 and SUZ12). Based on this degradation preference for BMI1 and RING1B over EED and the fact that genetic depletion of EED did not lead to loss of BMI1 and RING1B, we conclude that the degradation of BMI1 and RING1B induced by MS147 is not a consequence of EED degradation. This successful example, which supports our novel protein complex degrader approach, could expand the pool of degradable proteins to include proteins that lack small-molecule binders, but interact with other proteins that have small-molecule binders. In addition, while EED is not a core component of PRC1 but interacts with BMI1 and RING1B, we demonstrated that it can still be exploited to bring BMI1 and RING1B to VHL, suggesting that it is possible to utilize a relatively weak binding partner protein to recruit the ubiquitination machinery to the target protein, thereby potentially further expanding the utility of this protein complex degrader approach. Lastly, it is not surprising that the linker difference in MS147 and PROTAC 2 contributes to their different degradation profiles, based on that isoform or mutant selective PROTACs have been achieved by utilizing appropriate linkers and/or E3 ligase ligands.^[18]^

PRC1, a prominent epigenetic protein complex, plays critical roles in cancer development and progression. However, chemical agents targeting PRC1 are extremely limited. This study provides a novel chemical tool, which selectively targets PRC1 over PRC2. To the best of our knowledge, MS147 is the first and only BMI1 and RING1B degrader, which selectively degrades PRC1 components, BMI1 and RING1B, over PRC2 components: EED, EZH2 and SUZ12, and effectively suppresses the proliferation in cancer cells that are insensitive to EZH2 knockout or EED/PRC2 degraders. This well-characterized chemical tool is valuable for the research community to further investigate the roles of PRC1 in physiology and pathophysiology. Furthermore, the novel protein complex degrader strategy presented in this study could potentially expand the pool of degradable protein targets.

## 4. Experimental Section

### Compound synthesis

Synthesis and characterization of compounds **1** – **10**, MS147N1, MS147N2, as well as synthetic intermediates, are described in the Supplementary Materials.

### Cell culture

DMEM medium with 10% FBS and 1% mixture of penicillin and streptomycin was used for K562, KARPAS-422 and NCI-H1299 cells. RPMI-1640 medium with 10% FBS and 1% mixture of penicillin and streptomycin was used for MDA-MB-231 cells.

### Western blotting

Compound-treated cells were collected and lysed using RIPA buffer (Thermo Fisher Scientific, USA) with protease and phosphatase inhibitor cocktail (Thermo Fisher Scientific, USA). After incubation for 30 min at 0 °C, samples were centrifuged for 15 min at 18000 rpm and 4°C. The supernatant was collected, mixed with Laemmli sample buffer (Bio-Rad, USA), and then heated at 100°C for 7 min. The Pierce Rapid Gold BCA kit (Thermo Fisher Scientific, USA) was used for protein quantification. Based on the result, 10 μg of each sample was used for SDS-PAGE and transferred on to PVDF using Trans-Blot Turbo Transfer system (Bio-Rad, USA). Membranes were blocked using PBS Odyssey Blocking Buffer (LI-COR, USA) for 1 h at room temperature, then incubated overnight at 4 °C with the following primary antibodies: EZH2 (5246S, Cell Signaling Technology), SUZ12 (3737S, Cell Signaling Technology), EED (85322S, Cell Signaling Technology), BMI1 (6964S, Cell Signaling Technology), RING1B (16031-1-AP, Proteintech), H3K27me3 (9733S, Cell Signaling Technology), H3 (4499L, Cell Signaling Technology), H2AK119ub (8240S, Cell Signaling Technology), H2A (12349S, Cell Signaling Technology), Vinculin (13901S, Cell Signaling Technology), VHL (68547S, Cell Signaling Technology). Blots were imaged using Odyssey system (LI-COR, USA) and quantified using Image Studio (LI-COR, USA).

### Preparation of EED and VHL proteins for ITC study

For the expression of EED C-terminal domain (75-441), pET28-GST-LIC-EED vector was obtained from Addgene (Plasmid number 25311). The plasmid was transformed into *Escherichia coli* BL21 (DE3) cells and grown in TB medium at 37 °C until the culture reached an OD600 of ∼ 2.8. After cooling down to 15 °C, protein expression was induced by IPTC (0.4 mM) followed by incubation for 16 h. The cell was harvested and resuspended in a lysis buffer (50 mM phosphate buffer pH 7.5, 1 M NaCl, 5% glycerol) with 0.2 mM AEBSF. After sonication to lyse, the lysate was clarified by centrifugation and applied onto GSTrap FF 5 mL (GE Healthcare, USA) using a buffer (50 mM Tris-HCl pH 7.5, 150 mM NaCl). After washing, the column was filled by syringe with 5 mL of thrombin solution (20 U/mL in PBS, pH 7.3) (Sigma-Aldrich, USA) and sealed to cleavage GST-Tag. The resulting column was incubated for 16 h at 4 °C and washed with a buffer (50 mM Tris-HCl pH 7.5, 150 mM NaCl) to collect the GST-tag cleaved protein. After confirming cleavage by SDS-PAGE, size-exclusion column (SEC) using HiLoad 26/600 Superdex 200 (GE Healthcare, USA) was performed to separate thrombin and the GST-tag cleaved protein in a SEC buffer (50 mM Tris-HCl pH 7.5, 150 mM NaCl). After concentration of purified protein, the protein was flash frozen and stored at -80 °C for further usage.

For the expression of His-tagged VCB complex, VHL (54-213) with N-terminal His tag and a TEV protease cleavable site and EloB (1-104) and EloC (1-112) (Q15369) in pCDF Duet vector were co-transformed into *Escherichia coli* BL21 (DE3) cells. The cells were grown in LB medium at 37 °C until the culture reached an OD600 of ∼ 0.8. After cooling down to 18 °C, protein expression was induced by IPTC (0.4 mM) followed by incubation for 16 h. The cell was harvested and resuspended in a lysis buffer (50 mM Tris-HCl pH 7.5, 500 mM NaCl, 5% glycerol, 0.01% IGEPAL, 25mM imidazole and 5mM β-ME) with Pierce Protease Inhibitor Tablets (Thermo Fisher Scientific, USA) and 1 mM PMSF. After sonication to lyse, the lysate was clarified by centrifugation and applied onto HisTrap 5 mL (GE Healthcare, USA) using an imidazole gradient from 25 – 250 mM. The protein was further purified by size exclusion column (SEC) using HiLoad 26/600 Superdex 200 (GE Healthcare, USA) in SEC buffer (50 mM Tris-HCl pH 7.5, 150 mM NaCl, 2 mM TCEP). After concentration of purified protein, the protein was flash frozen and stored at -80 °C for further usage.

### Isothermal titration calorimetry (ITC)

ITC experiments were performed to assess binding affinities of MS147 and its negative controls (MS147N1 and MS147N2) for EED (75-441) and VCB complex using a MicroCal iTC200 (Malvern, UK). For determining binding affinity to EED, 13 injections from the syringe solution (400 μM of EED) were titrated into 300 μL of the cell solution (40 μM of compounds) with stirring at 750 rpm in 20 mM HEPES pH 7.4, 150 mM NaCl, 1% DMSO. For determining binding affinity to VCB, 19 injections from the syringe solution (200 μM of compounds) were titrated into 300 μL of the cell solution (40 μM of VCB complex) with stirring at 750 rpm in 25 mM Tris-HCl pH 7.5, 500 mM NaCl, 5% DMSO. The data were fitted by single binding site model using Microcal Origin 7.0 (Malvern). The reported values represent the mean ± SD from two independent experiments.

### shRNA-mediated EED knockdown experiment in K562 cells

pLKO.1 short hairpin RNA (shRNA) vector targeting human EED (TRCN0000021205) was purchased from MilliporeSigma. To generate an EED knockdown stable cell line, 293T cells were seeded on 100 mm tissue culture dishes at a density of 4 × 10^6^ cells per dish. The next day, cells were transfected with lentiviral vector and lentiviral packaging plasmids including psPAX2 and pMD2.G using PEI transfection reagent (MilliporeSigma). Virus was harvested at 48 hours post transfection, filtered through a 0.45μm syringe filter, and diluted in a complete medium supplemented with 5 μg/ml polybrene (MilliporeSigma) to infect K562 cells. Using the generated virus, K562 cells were infected for 24 hours in the presence of puromycin as 2 μg/mL. Then, MS147 was treated for an additional 24 h at 5 μM with DMSO as control. The protein levels of EED, BMI1 and RING1B with vinculin as a control were monitored by Western blot. The data shown in the figure are representative of two independent experiments.

### Quantitative reverse transcriptase polymerase chain reaction (RT-qPCR)

The real-time quantitative polymerase chain reaction (RT-qPCR) was performed as described previously.^[32]^ Briefly, KARPAS-422 cells were treated with either DMSO or MS147 at 5 μM for 24 h in 6-well plate. Subsequently, the samples were pelleted by centrifugation at 15,000 rpm for 10 min at 4°C. Total RNA was extracted using the Monarch® Total RNA Miniprep Kit (T2010S, New England Biolabs), and cDNA was generated using the SuperScript™ IV First-Strand Synthesis System (18091050, Thermo Fisher). qPCR was performed using the Agilent Technologies Stratagene Mx3005p qPCR system. The primer sets used for RT-PCR are listed below:

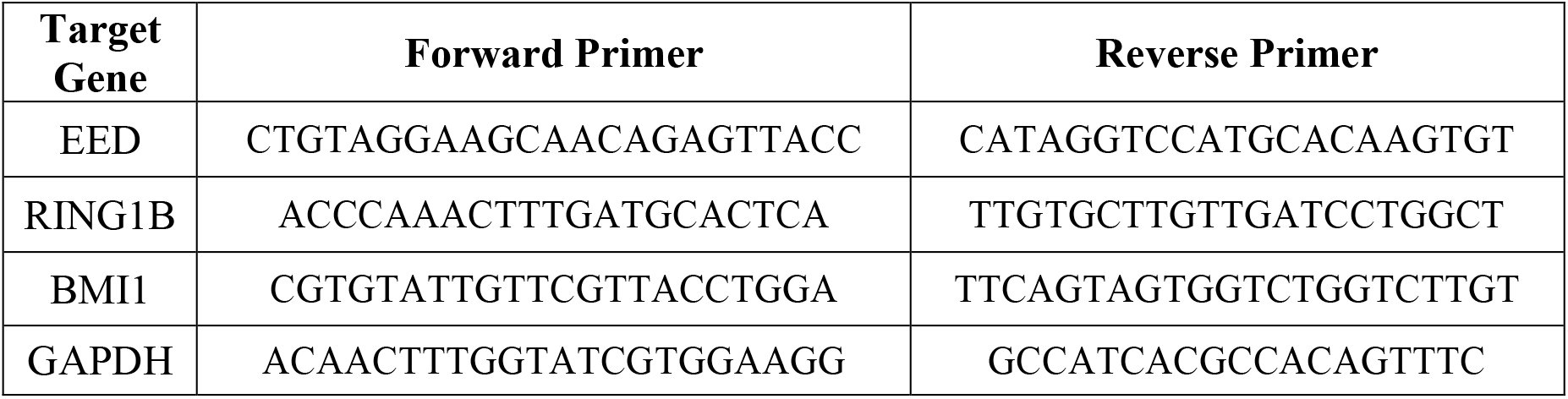

### siRNA-mediated EED knockdown (KD) experiment in MDA-MB-231 cells

Validated Silencer® Select pre-designed siRNA targeting EED was purchased from Thermo Fisher Scientific. MDA-MB-231 cells were seeded at a density of 500,000 cells/well in a 6-well plate and transfected with siRNA targeting EED using Lipofectamine RNAiMAX Transfection Reagent (Thermo Fisher Scientific) for 2 days, according to instruction. The KD efficiency was analyzed by Western blot. After confirming KD efficiency, the protein levels of EED, EZH2 and SUZ12 with vinculin as a control were monitored by Western blot. The data shown in the figure are representative of two independent experiments.

### Immunoprecipitation

K562 cells were seeded as 2 ×10^6^ in 10 cm dish. After 24 h, cells were harvested and washed with ice-cold PBS twice. The pellet was lysed with 500 μL of RIPA (Thermo Fisher Scientific, USA) with EDTA-free phosphatase and protease inhibitor (Thermo Fisher Scientific, USA) and 2 mM TCEP. After centrifugation to remove cell debris, the lysate was applied on Pierce Co-Immunoprecipitation kit (Thermo Fisher Scientific, USA) according to instructions from company with followed modification. Cell lysate was pre-incubated with VHL antibody (68547S, Cell Signaling Technology), and then added to agarose resin complex, where compounds (DMSO, MS147, MS147N1) were treated for 3 h as 40 μM. The resin was washed and eluted by sample buffer and then, analyzed by Western blotting. The data shown in the figure are representative of two independent experiments.

### Selectivity assays

Selectivity assays against other methyltransferases and reader domains were performed by Reaction Biology Corp. (USA) using the ^3^H-labeled SAM based assay for methyltransferases and thermal shift assay for reader domains. All of experiments were performed using 10 μM of MS147 in duplicate. The compound enzyme inhibition effect was calculated as percentage of inhibition against control enzyme activity.

### Cell viability assay

K562, MDA-MB-231, NCI-H1299 and PNT2 cells were seeded in 96-well plates (Thermo Fisher Scientific, USA) at 1×10^4^ cells per well and treated with DMSO or the indicated compound (EED226, PROTAC 2 or MS147) for 5 days at various concentrations. Cell viability was evaluated using CCK-8 (Dojindo, USA) following its protocol. All of values were plotted using GraphPad Prism 8. The data shown in the figures represent the means ± SD from three independent experiments.

### Statistical analysis

The Western blot data shown in the figure are representative of two biological independent experiments. The statistical analysis of cell viability assay, selectivity assay and RT-qPCR experiments were conducted using Prism 8 (GraphPad Software, USA), and shown as means ± SD from two or three independent experiments.

## Acknowledgements

This research was supported in part by an endowed professorship by the Icahn School of Medicine at Mount Sinai (to J.J.). This work utilized the NMR Spectrometer Systems at Mount Sinai acquired with funding from the U.S. National Institutes of Health (NIH) SIG grants 1S10OD025132 and 1S10OD028504. J.J. acknowledges the support by the grants R01CA218600, R01CA230854, R01CA260666, R01CA268384 and R01CA268519 from the National Cancer Institute (NCI) at the NIH. M.K. acknowledges the support by the Training Grant in Cancer Biology (T32CA078207) from the NCI. Y.Z. acknowledges the support by the National Institute of General Medical Sciences (NIGMS)-funded Integrated Pharmacological Sciences Training Program T32GM062754. B.D. acknowledges the support from the Medical Scientist Training Program (training grant T32GM007280) at the Icahn School of Medicine at Mount Sinai and the grant (3R01CA230854S1) from the NCI.

## Conflict of interests

J.J. is a cofounder and equity shareholder in Cullgen Inc. and a consultant for Cullgen Inc., EpiCypher Inc., and Accent Therapeutics Inc. The Jin laboratory received research funds from Celgene Corporation, Levo Therapeutics, Cullgen, Inc., and Cullinan Oncology.

## Contributions

K. –S. P., L.Q., M.K. and J.J. conceived the project and designed experiments. L.Q. and K.L. synthesized all compounds. K. –S. P. and M.K. performed western blot experiments and cell viability experiments. K. –S. P. and B.D. performed ITC experiments. M.K. performed RT-qPCR experiments. K. –S. P., A.R.K. and G.G.W. designed and performed EED KD experiments. All authors discussed the results and commented on the manuscript. K. –S. P., L.Q., M.K., H.Ü.K., Y.Z. and J.J. analyzed the data. K. –S. P., H.Ü.K., Y.Z. and J.J. wrote the paper. K. –S. P., L.Q. and M.K. contributed equally to this work.

