## Supporting Information for "Targeted degradation of PRC1 components, BMI1 and RING1B, via a novel protein complex degrader strategy"

**Corresponding Authors**

### Table of contents

**Supplemental Figures S1-S7**

**Figure S1.** Binding between EED and PRC1 components, BMI1 and RING1B, confirmed using immunoprecipitation.

**Figure S2.** Design of putative degraders of BMI1 and RING1B.

**Figure S3.** The effect of compounds **1-10** on reducing the protein levels of BMI1, RING1B, EED and H2AK119ub.

**Figure S4.** MOA of PROTAC 2-mediated degradation of PRC2 components, EED, EZH2 and SUZ12.

**Figure S5.** EZH2 and SUZ12 are destabilized following EED KD in MDA-MB-231 cells.

**Figure S6.** MS147 degrades RING1A in K562 cells.

**Figure S7.** MS147, EED226 and PROTAC 2 are non-toxic in PNT2 cells, a normal human prostate cell line.

**Synthetic procedures and chemical characterization data for all new compounds**

**$^{1}$H NMR and $^{13}$C NMR spectra of key compounds**

**References**

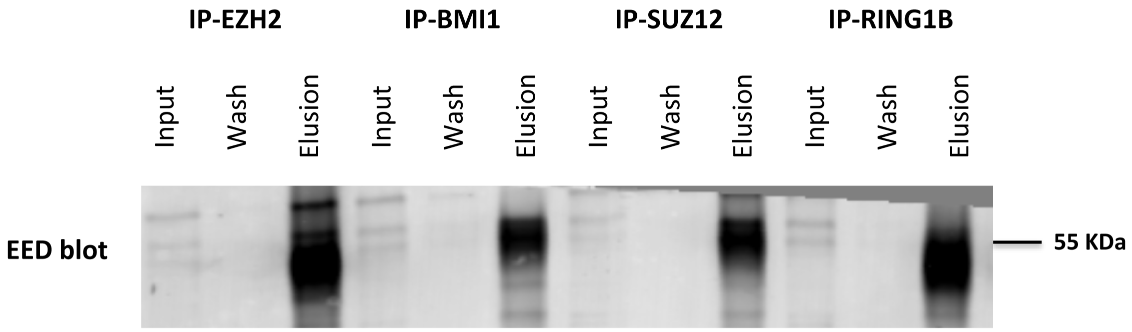


**Figure S1.** Binding between EED and PRC1 components, BMI1 and RING1B, confirmed using immunoprecipitation. Co-elusion of EED was confirmed by Western blot in K562 cells after IP with BMI1 or RING1B antibody. EZH2 and SUZ12 were used as positive controls.

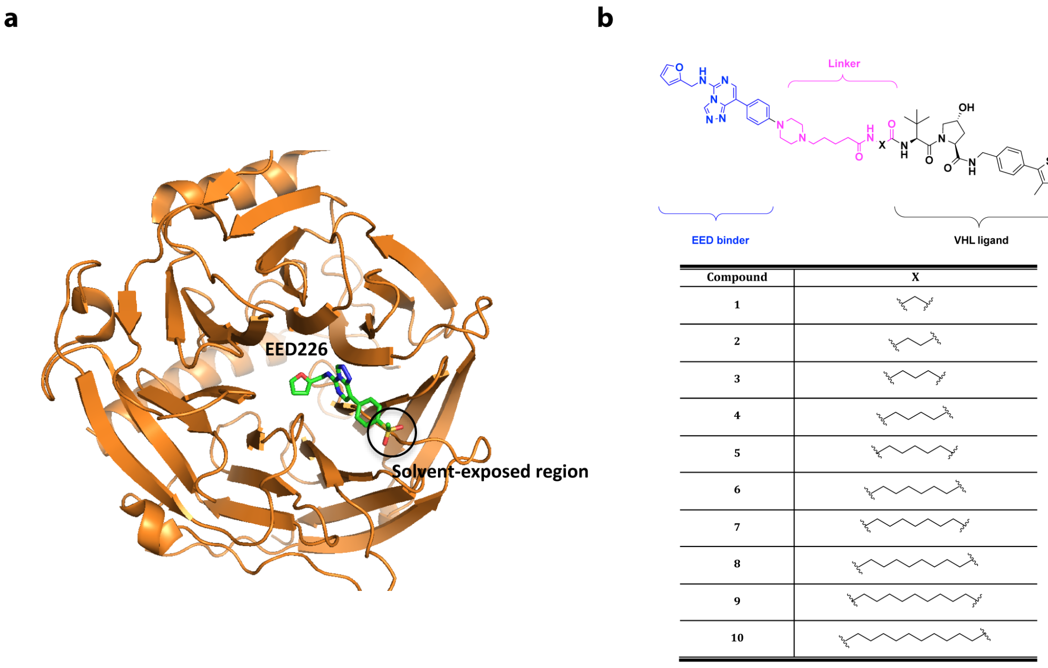


**Figure S2**. Design of putative degraders of BMI1 and RING1B. (a) Co-crystal structure of EED in complex with EED226 (PDB: 5WUK). The black circle indicates a solvent-exposed region of EED226, which could be used to attach a linker. (b) Chemical structures of the designed BMI1 and RING1B degraders **1** – **10**.

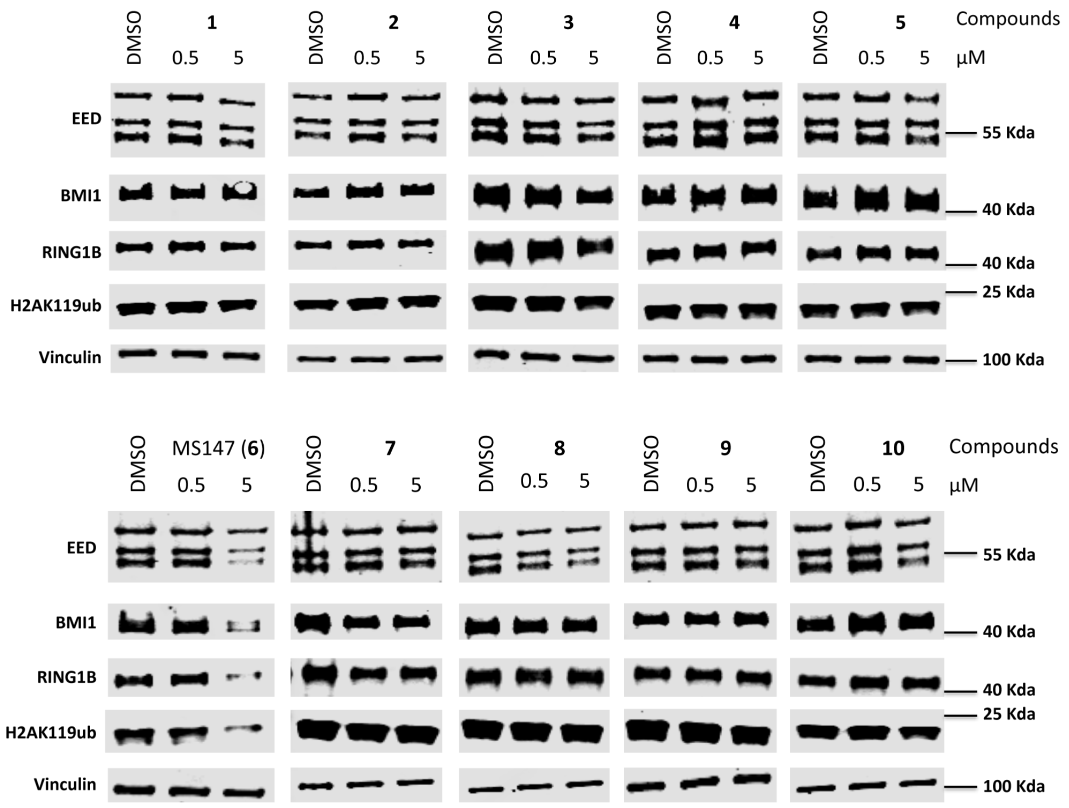


**Figure S3**. The effect of compounds **1-10** on reducing the protein levels of BMI1, RING1B, EED and H2AK119ub. K562 cells were treated with the indicated compound at the indicated concentrations for 24 h. The protein levels of BMI1, RING1B, EED and H2AK119ub were determined using WB with vinculin as the loading control. The results shown are representative of two independent experiments.

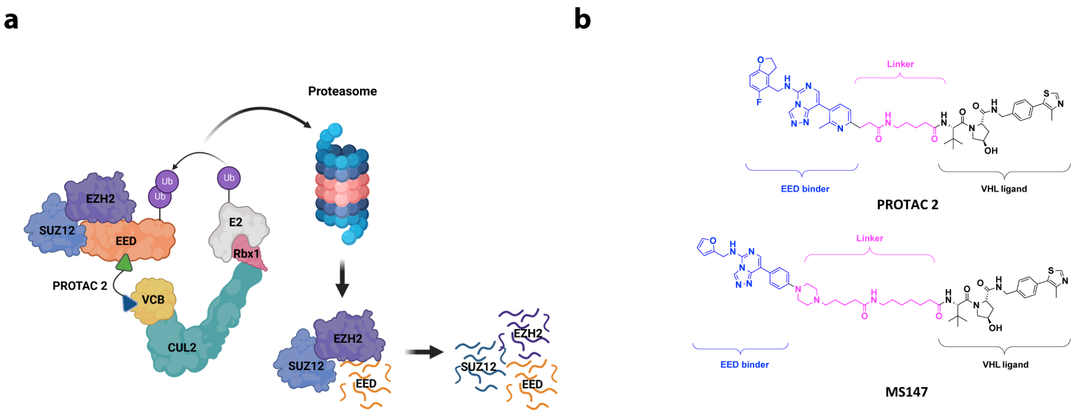


**Figure S4**. MOA of PROTAC 2-mediated degradation of PRC2 components, EED, EZH2 and SUZ12. (a) Illustration of the MOA of PROTAC 2 for degrading EED first and other PRC2 components (EZH2 and SUZ12) subsequently based on previously published results[1]. (b) Chemical structure of PROTAC 2 in comparison to that of MS147.

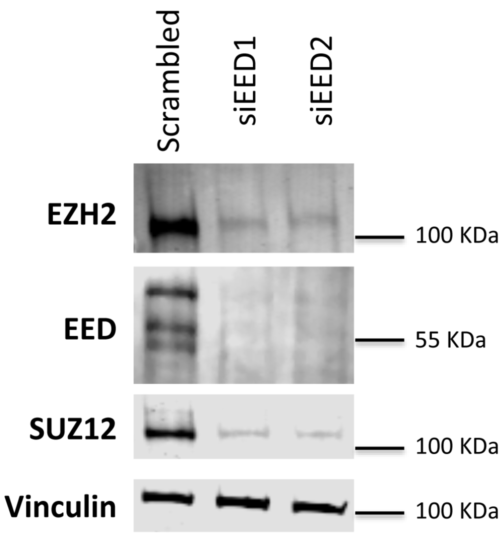


**Figure S5**. EZH2 and SUZ12 are destabilized following EED KD in MDA-MB-231 cells. MDA-MB-231 cells were transfected via lipofectamine with siEED for 48 h. After EED KD, the protein levels of EZH2 and SUZ12 in addition to EED were determined using WB with vinculin as the loading control. The results shown are representative of two independent experiments.

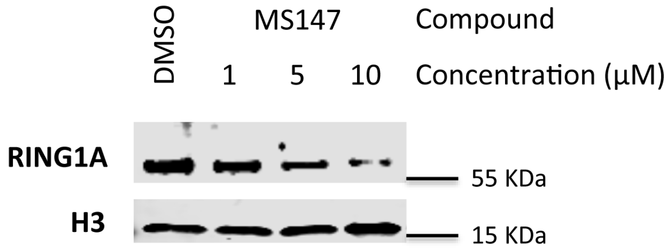


**Figure S6**. MS147 degrades RING1A in K562 cells. K562 cells were treated with MS147 at the indicated concentrations for 24 h. The protein level of RING1A was determined using WB with H3 as the loading control. The results shown are representative of two independent experiments. .

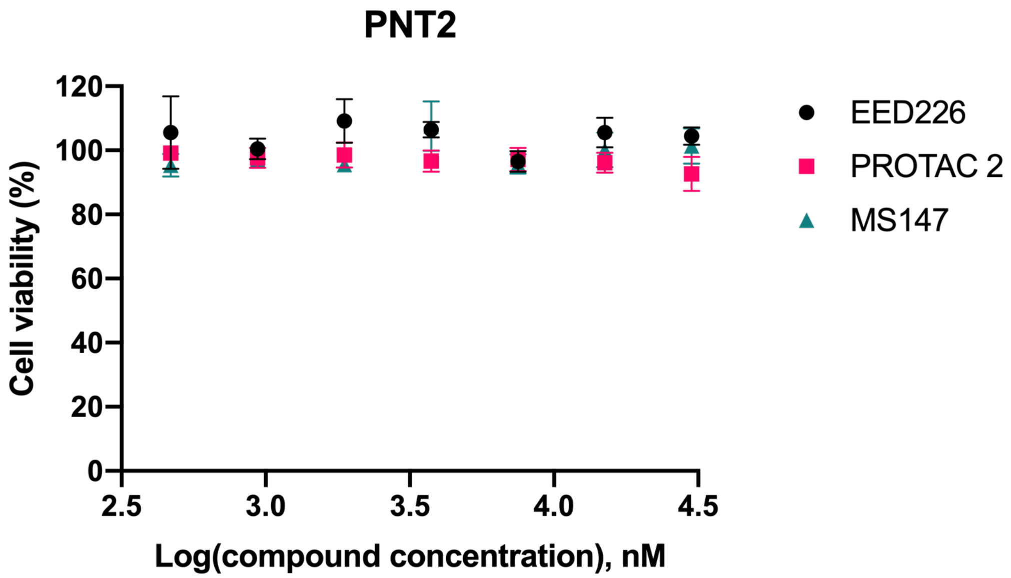


**Figure S7**. MS147, EED226 and PROTAC 2 are non-toxic in PNT2 cells, a normal human prostate cell line. PNT2 cells were treated with serial dilution of EED226, PROTAC 2 or MS147 for 5 days. Cell viability was monitored by the CCK-8 assay. The data shown represent the means ± SD from three independent experiments.

**Synthetic procedures and chemical characterization data for all new compounds**

**General chemistry procedures**

All chemical reagents were purchased from commercial vendors and used without further purification. The flash column chromatography was conducted using a Teledyne ISCO CombiFlash Rf+ instrument. This instrument was also equipped with a variable-wavelength UV detector and a fraction collector. RediSep Rf Gold C18 columns were used for purification. High-performance liquid chromatography (HPLC) spectra for compounds were acquired using an Agilent 1200 Series system with a DAD detector. Chromatography was performed on a 2.1 × 150 mm Zorbax 300SB-C18 5 μm column with water containing 0.1% formic acid as solvent A and acetonitrile containing 0.1% formic acid as solvent B at a flow rate of 0.4 mL/min. The gradient program was as follows: 1% B (0−1 min), 1−99% B (1−4 min), and 99% B (4−8 min). Ultra performance liquid chromatography (UPLC) spectra for compounds were acquired using a Waters Acquity I-Class UPLC system with a PDA detector. Chromatography was performed on a 2.1 × 30 mm ACQUITY UPLC BEH C18 1.7 μm column with water containing 3% acetonitrile, 0.1% formic acid as solvent A and acetonitrile containing 0.1% formic acid as solvent B at a flow rate of 0.8 mL/min. The gradient program was as follows: 1−99% B (1−1.5 min), and 99−1% B (1.5−2.5 min). High-resolution mass spectra (HRMS) data were acquired in the positive ion mode using Agilent G1969A API-TOF with an electrospray ionization (ESI) source. Nuclear magnetic resonance (NMR) spectra were acquired on a Bruker DRX-500 spectrometer with 500 MHz for proton ($^{1}H$ NMR) or a Bruker DRX-600 spectrometer with 600 MHz for proton ($^{1}H$ NMR) or 151 MHz for carbon ($^{13}C$ NMR); Chemical shifts are reported in ppm (δ). Preparative HPLC was performed using an Agilent Prep 1200 series with UV detector set to 220 nm. Samples were injected into a Phenomenex Luna 75 × 30 mm, 5 µm, C18 column at room temperature. The flow rate was 40 mL/min. A linear gradient was used with 10% of Acetonitrile (A) in $H_2O$ (with 0.1% TFA) (B) to 100% of Acetonitrile (A). All final compounds had > 95% purity using the UPLC and HPLC methods described above.

**Scheme S1. The synthesis of Compounds 1-10 and MS147N1**

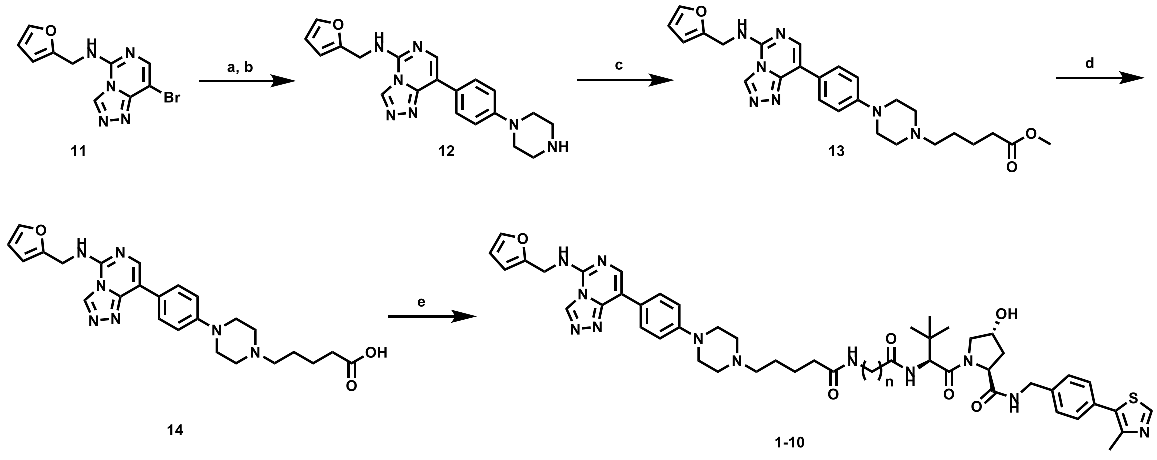


Reaction Condition: (a) 4-(4-*tert* Butoxycarbonylpiperazinyl)phenylboronic acid pinacol ester, Pd(dppf)$Cl_2$, $K_2CO_3$, dioxane, $H_2O$, 130 °C; (b) TFA, DCM, rt; (c) methyl 5-bromovalerate, $K_2CO_3$, DIEA, DMF, rt; (d) LiOH, MeOH, $H_2O$, THF; (e) EDCI, HOAt, NMM, DMSO, rt.

##### *N*-(furan-2-ylmethyl)-8-(4-(piperazin-1-yl)phenyl)-[1,2,4]triazolo[4,3-*c*]pyrimidin-5-amine (12)

A solution of **11** [2] (500 mg, 1.7 mmol), 4-(4-*tert* Butoxycarbonylpiperazinyl)-phenylboronic acid pinacol ester (660 mg, 1.7 mmol), Pd(dppf)$Cl_2$ (14 mg, 0.17 mmol), potassium carbonate (234 mg, 1.7 mmol) in dioxane and water was heated to 130 °C under microwave for 1 h. After cooling to room temperature, the mixture was poured in water and extracted with EtOAc. The combined organic layer was washed with brine and concentrated *in vacuo*. The resulting residue was purified by silica gel flash chromatography to give the compound as yellow solid (750 mg, 92%). The obtained intermediate was dissolved in 5 mL DCM, to the resulting solution was added 3 mL TFA. After being stirred for 1 h at room temperature, the reaction mixture was concentrated and the residue was purified by reverse phase C18 column (10% - 100% methanol / 0.1% TFA in water) to afford the compound **12** as a white solid in TFA salt form (600 mg, 80%). $^1$H NMR (600 MHz, Methanol-$d_4$) δ 8.42 (s, 1H), 8.12 (s, 1H), 7.87 – 7.83 (m, 2H), 7.46 (t, $J$ = 1.4 Hz, 1H), 7.17 – 7.14 (m, 2H), 6.39 – 6.37 (m, 2H), 4.85 (s, 2H), 3.53 – 3.49 (m, 4H), 3.44 – 3.40 (m, 4H). MS (ESI) $m/z$ = 376.2 $[M + H]^+$.

**Methyl 5-(4-(4-(5-((furan-2-ylmethyl)amino)-[1,2,4]triazolo[4,3-*c*]pyrimidin-8-yl)phenyl)piperazin-1-yl)pentanoate (13)**

A solution of **12** (600 mg, 1.22 mmol) and methyl 5-bromovalerate (238 mg, 1.22 mmol) in 10 mL of DMF was treated with $K_2CO_3$ (276 mg, 2 mmol) and DIEA (190 mg, 1.9 mmol). The resulting mixture was stirred at RT overnight. After the reaction was completed, the reaction mixture was poured into ice water, aqueous phase was extracted with ethyl acetate. The combined organic phase was washed with brine twice, dried and concentrated. The resulting residue was used directly for the next step without further purification. MS (ESI) $m/z$ = 490.3 $[M + H]^+$.

**5-(4-(4-(5-((Furan-2-ylmethyl)amino)-[1,2,4]triazolo[4,3-*c*]pyrimidin-8-yl)phenyl)piperazin-1-yl)pentanoic acid (14)**

To a solution of the obtained intermediate **13** (450 mg, 0.92mmol) in 5 mL MeOH, 5 mL $H_2O$, and 5 mL THF, LiOH (30 mg, 1 mmol) was added. The mixture was stirred at RT overnight. Then the mixture was purified by reverse phase C18 column (10% - 100% methanol / 0.1% TFA in water) to afford the compound **14** as a white solid in TFA salt form (482 mg, 89%). $^1$H NMR (600 MHz, Methanol-$d_4$) δ 8.38 (s, 1H), 8.05 (s, 1H), 7.79 – 7.73 (m, 2H), 7.44 – 7.41 (m, 1H), 7.07 – 7.02 (m, 2H), 6.36 – 6.33 (m, 2H), 4.79 (s, 2H), 3.87 (s, 2H), 3.71 – 3.61 (m, 2H), 3.26 – 3.16 (m, 4H), 3.14 – 3.06 (m, 2H), 2.41 (t, *J* = 7.1 Hz, 2H), 1.88 – 1.79 (m, 2H), 1.69 (p, *J* = 7.3 Hz, 2H). MS (ESI) *m/z* = 476.3 $[M + H]^+$.

**(2*S*,4*R*)-1-((*S*)-2-(2-(5-(4-(4-(5-((furan-2-ylmethyl)amino)-[1,2,4]triazolo[4,3-*c*]pyrimidin-8-yl)phenyl)piperazin-1-yl)pentanamido)acetamido)-3,3-dimethylbutanoyl)-4-hydroxy-*N*-(4-(4-methylthiazol-5-yl)benzyl)pyrrolidine-2-carboxamide (1)**

To a solution of compound **14** (12 mg, 0.02 mmol) in DMSO (1 mL), was added (2*S*,4*R*)-1-((*S*)-2-(2-aminoacetamido)-3,3-dimethylbutanoyl)-4-hydroxy-*N*-(4-(4-methylthiazol-5-yl)benzyl)pyrrolidine-2-carboxamide (14.3 mg, 0.02 mmol, 1.0 equiv), EDCI (5.8 mg, 0.03 mmol, 1.5 equiv), HOAt (4.1 mg, 0.03 mmol, 1.5 equiv), and NMM (6.1 mg, 0.06 mmol, 3.0 equiv). The resulting mixture was stirred at room temperature overnight and then purified by preparative HPLC to give the title compound as a white solid in TFA salt form (15.7 mg, 74%). $^1$H NMR (600 MHz, Methanol-$d_4$) δ 9.07 (s, 1H), 8.43 (s, 1H), 8.10 (s, 1H), 7.83 (d, *J* = 8.5 Hz, 2H), 7.48 – 7.40 (m, 5H), 7.15 (d, *J* = 8.4 Hz, 2H), 6.39

– 6.36 (m, 2H), 4.84 (s, 2H), 4.68 – 4.64 (m, 1H), 4.59 – 4.48 (m, 3H), 4.38 (d, *J* = 15.5 Hz, 1H), 3.99 – 3.88 (m, 5H), 3.81 (dd, *J* = 10.9, 3.9 Hz, 1H), 3.74 – 3.68 (m, 2H), 3.30 – 3.24 (m, 4H), 3.14 (t, *J* = 12.6 Hz, 2H), 2.49 (s, 3H), 2.41 (t, *J* = 6.8 Hz, 2H), 2.27 – 2.22 (m, 1H), 2.12 – 2.06 (m, 1H), 1.92 – 1.85 (m, 2H), 1.81 – 1.75 (m, 2H), 1.06 (s, 9H). HRMS *m/z* $[M + H]^+$ calcd for $C_{49}H_{61}N_{12}O_6S^+$ 945.4552, found 945.4563.

**(2*S*,4*R*)-1-((*S*)-2-(3-(5-(4-(4-(5-((furan-2-ylmethyl)amino)-[1,2,4]triazolo[4,3-*c*]pyrimidin-8-yl)phenyl)piperazin-1-yl)pentanamido)propanamido)-3,3-dimethylbutanoyl)-4-hydroxy-*N*-(4-(4-methylthiazol-5-yl)benzyl)pyrrolidine-2-carboxamide (2)**

Using the same procedure for the synthesis of compound **1** with (2*S*,4*R*)-1-((*S*)-2-(3-aminopropanamido)-3,3-dimethylbutanoyl)-4-hydroxy-*N*-(4-(4-methylthiazol-5-yl)benzyl)pyrrolidine-2-carboxamide to get compound **2** as a white solid in TFA salt form (14.7 mg, 68%). $^1$H NMR (600 MHz, Methanol-$d_4$) δ 8.99 (s, 1H), 8.43 (s, 1H), 8.10 (s, 1H), 7.83 (d, *J* = 8.5 Hz, 2H), 7.50 – 7.44 (m, 3H), 7.41 (d, *J* = 8.0 Hz, 2H), 7.13 (d, *J* = 8.6 Hz, 2H), 6.39 – 6.36 (m, 2H), 4.84 (s, 2H), 4.64 (s, 1H), 4.59 – 4.49 (m, 3H), 4.43 – 4.37 (m, 1H), 4.00 – 3.91 (m, 3H), 3.83 (dd, *J* = 10.9, 3.8 Hz, 1H), 3.74 – 3.68 (m, 2H), 3.54 – 3.47 (m, 1H), 3.46 – 3.40 (m, 1H), 3.30 – 3.20 (m, 4H), 3.13 (t, *J* = 12.6 Hz, 2H), 2.53 – 2.49 (m, 2H), 2.47 (s, 3H), 2.32 – 2.25 (m, 3H), 2.14 – 2.08 (m, 1H), 1.85 – 1.78 (m, 2H), 1.75 – 1.68 (m, 2H), 1.06 (s, 9H). HRMS *m/z* $[M + H]^+$ calcd for $C_{50}H_{63}N_{12}O_6S^+$ 959.4709, found 959.4707.

**(2*S*,4*R*)-1-((*S*)-2-(4-(5-(4-(4-(5-((furan-2-ylmethyl)amino)-[1,2,4]triazolo[4,3-*c*]pyrimidin-8-yl)phenyl)piperazin-1-yl)pentanamido)butanamido)-3,3-dimethylbutanoyl)-4-hydroxy-*N*-(4-(4-methylthiazol-5-yl)benzyl)pyrrolidine-2-carboxamide (3)**

Using the same procedure for the synthesis of compound **1** with (2*S*,4*R*)-1-((*S*)-2-(4-aminobutanamido)-3,3-dimethylbutanoyl)-4-hydroxy-*N*-(4-(4-methylthiazol-5-yl)benzyl)pyrrolidine-2-carboxamide to get compound **3** as a white solid in TFA salt form (15.8 mg, 73%). $^{1}$H NMR (600 MHz, Methanol-$d_4$) δ 8.90 (s, 1H), 8.42 (s, 1H), 8.11 (s, 1H), 7.85 (d, *J* = 8.8 Hz, 2H), 7.47 – 7.44 (m, 3H), 7.43 – 7.40 (m, 2H), 7.16 (d, *J* = 8.9 Hz, 2H), 6.39 – 6.37 (m, 2H), 4.85 (s, 2H), 4.64 (s, 1H), 4.59 – 4.50 (m, 4H), 4.00 – 3.91 (m, 3H), 3.84 – 3.81 (m, 1H), 3.75 – 3.63 (m, 4H), 3.28 – 3.20 (m, 6H), 3.17 – 3.10 (m, 2H), 2.48 (s, 3H), 2.35 – 2.31 (m, 4H), 1.85 – 1.80 (m, 4H), 1.77 – 1.73 (m, 2H), 1.06 (s, 9H). HRMS *m/z* $[M + H]^+$ calcd for $C_{51}H_{65}N_{12}O_6S^+$ 973.4865, found 973.4863.

**(2*S*,4*R*)-1-((*S*)-2-(5-(5-(4-(4-(5-((furan-2-ylmethyl)amino)-[1,2,4]triazolo[4,3-*c*]pyrimidin-8-yl)phenyl)piperazin-1-yl)pentanamido)pentanamido)-3,3-dimethylbutanoyl)-4-hydroxy-*N*-(4-(4-methylthiazol-5-yl)benzyl)pyrrolidine-2-carboxamide (4)**

Using the same procedure for the synthesis of compound **1** with (2*S*,4*R*)-1-((*S*)-2-(5-aminopentanamido)-3,3-dimethylbutanoyl)-4-hydroxy-*N*-(4-(4-methylthiazol-5-yl)benzyl)pyrrolidine-2-carboxamide to get compound **4** as a white solid in TFA salt form (15.2 mg, 69%). $^{1}$H NMR (600 MHz, Methanol-$d_4$) δ 8.98 (s, 1H), 8.42 (s, 1H), 8.10 (s, 1H), 7.86 – 7.82 (m, 2H), 7.48 – 7.44 (m, 3H), 7.41 (d, *J* = 8.1 Hz, 2H), 7.15 (d, *J*

= 8.8 Hz, 2H), 6.39 – 6.36 (m, 2H), 4.84 (s, 2H), 4.65 – 4.62 (m, 1H), 4.59 – 4.50 (m, 3H), 4.37 (d, *J* = 15.5 Hz, 1H), 3.99 – 3.89 (m, 3H), 3.81 (dd, *J* = 10.9, 3.9 Hz, 1H), 3.71 (d, *J* = 12.2 Hz, 2H), 3.30 – 3.21 (m, 6H), 3.14 (t, *J* = 12.6 Hz, 2H), 2.48 (s, 3H), 2.35 – 2.30 (m, 4H), 2.26 – 2.20 (m, 1H), 2.12 – 2.07 (m, 1H), 1.87 – 1.80 (m, 2H), 1.76 – 1.71 (m, 2H), 1.67 – 1.62 (m, 2H), 1.58 – 1.52 (m, 2H), 1.05 (s, 9H). HRMS *m/z* $[M + H]^+$ calcd for $C_{52}H_{67}N_{12}O_6S^+$ 987.5022, found 987.5002.

**(2*S*,4*R*)-1-((*S*)-2-(6-(5-(4-(4-(5-((furan-2-ylmethyl)amino)-[1,2,4]triazolo[4,3-*c*]pyrimidin-8-yl)phenyl)piperazin-1-yl)pentanamido)hexanamido)-3,3-dimethylbutanoyl)-4-hydroxy-*N*-(4-(4-methylthiazol-5-yl)benzyl)pyrrolidine-2-carboxamide (5)**

Using the same procedure for the synthesis of compound **1** with (2*S*,4*R*)-1-((*S*)-2-(6-aminohexanamido)-3,3-dimethylbutanoyl)-4-hydroxy-*N*-(4-(4-methylthiazol-5-yl)benzyl)pyrrolidine-2-carboxamide to get compound **5** as a white solid in TFA salt form (14.3 mg, 64%). $^1$H NMR (600 MHz, Methanol-$d_4$) δ 9.00 (s, 1H), 8.42 (s, 1H), 8.11 (s, 1H), 7.84 (d, *J* = 8.7 Hz, 2H), 7.49 – 7.41 (m, 5H), 7.15 (d, *J* = 8.8 Hz, 2H), 6.39 – 6.36 (m, 2H), 4.84 (s, 2H), 4.65 (s, 1H), 4.60 – 4.50 (m, 3H), 4.38 (d, *J* = 15.5 Hz, 1H), 3.99 – 3.90 (m, 3H), 3.82 (dd, *J* = 10.9, 3.9 Hz, 1H), 3.71 (d, *J* = 12.0 Hz, 2H), 3.29 – 3.09 (m, 8H), 2.49 (s, 3H), 2.34 – 2.21 (m, 5H), 2.13 – 2.07 (m, 1H), 1.86 – 1.80 (m, 2H), 1.75 – 1.71 (m, 2H), 1.68 – 1.61 (m, 2H), 1.58 – 1.51 (m, 2H), 1.41 – 1.35 (m, 2H), 1.05 (s, 9H). HRMS *m/z* $[M + H]^+$ calcd for $C_{53}H_{69}N_{12}O_6S^+$ 1001.5178, found 1001.5177.

**(2*S*,4*R*)-1-((*S*)-2-(7-(5-(4-(4-(5-((furan-2-ylmethyl)amino)-[1,2,4]triazolo[4,3-*c*]pyrimidin-8-yl)phenyl)piperazin-1-yl)pentanamido)heptanamido)-3,3-dimethylbutanoyl)-4-hydroxy-*N*-(4-(4-methylthiazol-5-yl)benzyl)pyrrolidine-2-carboxamide (6, MS147)**

Using the same procedure for the synthesis of compound **1** with (2*S*,4*R*)-1-((*S*)-2-(7-aminoheptanamido)-3,3-dimethylbutanoyl)-4-hydroxy-*N*-(4-(4-methylthiazol-5-yl)benzyl)pyrrolidine-2-carboxamide to get the compound **MS147** as a white solid in TFA salt form (16.2 mg, 72%). $^{1}$H NMR (600 MHz, Methanol-$d_4$) δ 9.03 (s, 1H), 8.43 (s, 1H), 8.11 (s, 1H), 7.85 (d, $J$ = 8.7 Hz, 2H), 7.50 – 7.42 (m, 5H), 7.15 (d, $J$ = 8.7 Hz, 2H), 6.39 – 6.37 (m, 2H), 4.84 (s, 2H), 4.65 (s, 1H), 4.60 – 4.50 (m, 3H), 4.38 (d, $J$ = 15.5 Hz, 1H), 4.00 – 3.90 (m, 3H), 3.82 (dd, $J$ = 11.0, 4.0 Hz, 1H), 3.74 – 3.68 (m, 2H), 3.31 – 3.23 (m, 4H), 3.20 (t, $J$ = 7.1 Hz, 2H), 3.13 (t, $J$ = 12.5 Hz, 2H), 2.50 (s, 3H), 2.34 – 2.21 (m, 5H), 2.12 – 2.07 (m, 1H), 1.86 – 1.80 (m, 2H), 1.75 – 1.69 (m, 2H), 1.66 – 1.59 (m, 2H), 1.56 – 1.50 (m, 2H), 1.40 – 1.35 (m, 4H), 1.05 (s, 9H). $^{13}$C NMR (151 MHz, Methanol-$d_4$) δ 174.5, 174.4, 173.1, 170.9, 153.2, 151.7, 151.5, 151.4, 150.8, 147.6, 145.5, 142.0, 141.9, 138.8, 132.0, 130.1, 128.9, 128.1, 127.5, 124.4, 115.7, 114.1, 110.0, 107.0, 69.7, 59.4, 57.9, 57.5, 56.6, 52.8, 48.4, 48.3, 42.3, 38.9, 37.5, 37.3, 35.5, 35.1, 29.7, 28.9, 28.5, 26.3, 25.6, 25.6, 25.5, 23.6, 14.4. HRMS *m/z* $[M + H]^+$ calcd for $C_{54}H_{71}N_{12}O_6S^+$ 1015.5335, found 1015.5353.

**(2*S*,4*R*)-1-((*S*)-2-(8-(5-(4-(4-(5-((furan-2-ylmethyl)amino)-[1,2,4]triazolo[4,3-*c*]pyrimidin-8-yl)phenyl)piperazin-1-yl)pentanamido)octanamido)-3,3-**

**dimethylbutanoyl)-4-hydroxy-*N*-(4-(4-methylthiazol-5-yl)benzyl)pyrrolidine-2-carboxamide (7)**

Using the same procedure for the synthesis of compound **1** with (2*S*,4*R*)-1-((*S*)-2-(8-aminooctanamido)-3,3-dimethylbutanoyl)-4-hydroxy-*N*-(4-(4-methylthiazol-5-yl)benzyl)pyrrolidine-2-carboxamide to get the compound **7** as a white solid in TFA salt form (15 mg, 66%). $^{1}$H NMR (600 MHz, Methanol-$d_4$) δ 9.00 (s, 1H), 8.43 (s, 1H), 8.11 (s, 1H), 7.85 (d, $J$ = 8.7 Hz, 2H), 7.50 – 7.42 (m, 5H), 7.15 (d, $J$ = 8.6 Hz, 2H), 6.39 – 6.36 (m, 2H), 4.84 (s, 2H), 4.66 (s, 1H), 4.60 – 4.50 (m, 3H), 4.40 – 4.35 (m, 1H), 4.00 – 3.90 (m, 3H), 3.82 (dd, $J$ = 10.9, 3.9 Hz, 1H), 3.70 (d, $J$ = 12.2 Hz, 2H), 3.31 – 3.23 (m, 4H), 3.19 (t, $J$ = 7.1 Hz, 2H), 3.16 – 3.09 (m, 2H), 2.49 (s, 3H), 2.34 – 2.21 (m, 5H), 2.13 – 2.07 (m, 1H), 1.85 – 1.80 (m, 2H), 1.76 – 1.70 (m, 2H), 1.65 – 1.59 (m, 2H), 1.55 – 1.49 (m, 2H), 1.40 – 1.34 (m, 6H), 1.05 (s, 9H). HRMS *m/z* $[M + H]^+$ calcd for $C_{55}H_{73}N_{12}O_6S^+$ 1029.5491, found 1029.5486.

**(2*S*,4*R*)-1-((*S*)-2-(9-(5-(4-(4-(5-((furan-2-ylmethyl)amino)-[1,2,4]triazolo[4,3-*c*]pyrimidin-8-yl)phenyl)piperazin-1-yl)pentanamido)nonanamido)-3,3-dimethylbutanoyl)-4-hydroxy-*N*-(4-(4-methylthiazol-5-yl)benzyl)pyrrolidine-2-carboxamide (8)**

Using the same procedure for the synthesis of compound **1** with (2*S*,4*R*)-1-((*S*)-2-(9-aminononanamido)-3,3-dimethylbutanoyl)-4-hydroxy-*N*-(4-(4-methylthiazol-5-yl)benzyl)pyrrolidine-2-carboxamide to get the compound **8** as a white solid in TFA salt form (16.1 mg, 69%). $^{1}$H NMR (600 MHz, Methanol-$d_4$) δ 8.99 (s, 1H), 8.43 (s, 1H), 8.12 (s, 1H), 7.85 (d, $J$ = 8.7 Hz, 2H), 7.50 – 7.41 (m, 5H), 7.15 (d, $J$ = 8.7 Hz, 2H), 6.39

– 6.36 (m, 2H), 4.84 (s, 2H), 4.65 (s, 1H), 4.60 – 4.50 (m, 3H), 4.38 (d, *J* = 15.5 Hz, 1H), 4.00 – 3.90 (m, 3H), 3.82 (dd, *J* = 11.1, 3.9 Hz, 1H), 3.71 (d, *J* = 12.1 Hz, 2H), 3.31 – 3.23 (m, 4H), 3.19 (t, *J* = 7.2 Hz, 2H), 3.13 (t, *J* = 12.9 Hz, 2H), 2.49 (s, 3H), 2.33 – 2.21 (m, 5H), 2.12 – 2.07 (m, 1H), 1.87 – 1.81 (m, 2H), 1.76 – 1.69 (m, 2H), 1.66 – 1.59 (m, 2H), 1.54 – 1.49 (m, 2H), 1.39 – 1.33 (m, 8H), 1.05 (s, 9H). HRMS *m/z* $[M + H]^+$ calcd for $C_{56}H_{75}N_{12}O_6S^+$ 1043.5648, found 1043.5677.

**(2*S*,4*R*)-1-((*S*)-2-(10-(5-(4-(4-(5-((furan-2-ylmethyl)amino)-[1,2,4]triazolo[4,3-*c*]pyrimidin-8-yl)phenyl)piperazin-1-yl)pentanamido)decanamido)-3,3-dimethylbutanoyl)-4-hydroxy-*N*-(4-(4-methylthiazol-5-yl)benzyl)pyrrolidine-2-carboxamide (9)**

Using the same procedure for the synthesis of compound **1** with (2*S*,4*R*)-1-((*S*)-2-(10-aminodecanamido)-3,3-dimethylbutanoyl)-4-hydroxy-*N*-(4-(4-methylthiazol-5-yl)benzyl)pyrrolidine-2-carboxamide to get the compound **9** as a white solid in TFA salt form (15.6 mg, 66%). $^1H$ NMR (600 MHz, Methanol-$d_4$) δ 8.96 (s, 1H), 8.42 (s, 1H), 8.11 (s, 1H), 7.85 (d, *J* = 8.8 Hz, 2H), 7.50 – 7.41 (m, 5H), 7.15 (d, *J* = 8.7 Hz, 2H), 6.39 – 6.36 (m, 2H), 4.84 (s, 2H), 4.65 (s, 1H), 4.61 – 4.50 (m, 3H), 4.37 (d, *J* = 15.5 Hz, 1H), 3.99 – 3.90 (m, 3H), 3.82 (dd, *J* = 11.0, 3.9 Hz, 1H), 3.73 – 3.68 (m, 2H), 3.30 – 3.23 (m, 4H), 3.19 (t, *J* = 7.2 Hz, 2H), 3.17 – 3.10 (m, 2H), 2.49 (s, 3H), 2.34 – 2.21 (m, 5H), 2.12 – 2.07 (m, 1H), 1.87 – 1.80 (m, 2H), 1.76 – 1.70 (m, 2H), 1.65 – 1.57 (m, 2H), 1.55 – 1.49 (m, 2H), 1.37 – 1.32 (m, 10H), 1.05 (s, 9H). HRMS *m/z* $[M + H]^+$ calcd for $C_{57}H_{77}N_{12}O_6S^+$ 1057.5804, found 1057.5825.

**(2*S*,4*R*)-1-((*S*)-2-(11-(5-(4-(4-(5-((furan-2-ylmethyl)amino)-[1,2,4]triazolo[4,3-*c*]pyrimidin-8-yl)phenyl)piperazin-1-yl)pentanamido)undecanamido)-3,3-dimethylbutanoyl)-4-hydroxy-*N*-(4-(4-methylthiazol-5-yl)benzyl)pyrrolidine-2-carboxamide (10)**

Using the same procedure for the synthesis of compound **1** with (2*S*,4*R*)-1-((*S*)-2-(11-aminoundecanamido)-3,3-dimethylbutanoyl)-4-hydroxy-*N*-(4-(4-methylthiazol-5-yl)benzyl)pyrrolidine-2-carboxamide to get the compound **10** as a white solid in TFA salt form (14.3 mg, 60%). $^1$H NMR (600 MHz, Methanol-$d_4$) δ 8.97 (s, 1H), 8.42 (s, 1H), 8.11 (s, 1H), 7.85 (d, *J* = 8.7 Hz, 2H), 7.49 – 7.41 (m, 5H), 7.15 (d, *J* = 8.7 Hz, 2H), 6.39 – 6.36 (m, 2H), 4.84 (s, 2H), 4.65 (s, 1H), 4.61 – 4.50 (m, 3H), 4.37 (d, *J* = 15.5 Hz, 1H), 3.99 – 3.89 (m, 3H), 3.82 (dd, *J* = 11.0, 3.9 Hz, 1H), 3.74 – 3.68 (m, 2H), 3.31 – 3.23 (m, 4H), 3.21 – 3.10 (m, 4H), 2.49 (s, 3H), 2.34 – 2.21 (m, 5H), 2.13 – 2.07 (m, 1H), 1.87 – 1.80 (m, 2H), 1.76 – 1.70 (m, 2H), 1.66 – 1.57 (m, 2H), 1.54 – 1.49 (m, 2H), 1.37 – 1.29 (m, 12H), 1.05 (s, 9H). HRMS *m/z* $[M + H]^+$ calcd for $C_{58}H_{79}N_{12}O_6S^+$ 1071.5961, found 1071.5977.

**(2*R*,4*S*)-1-((*S*)-2-(7-(5-(4-(4-(5-((furan-2-ylmethyl)amino)-[1,2,4]triazolo[4,3-*c*]pyrimidin-8-yl)phenyl)piperazin-1-yl)pentanamido)heptanamido)-3,3-dimethylbutanoyl)-4-hydroxy-*N*-(4-(4-methylthiazol-5-yl)benzyl)pyrrolidine-2-carboxamide (MS147N1)**

Using the same procedure for the synthesis of compound **1** with (2*R*,4*S*)-1-((*S*)-2-(7-aminoheptanamido)-3,3-dimethylbutanoyl)-4-hydroxy-*N*-(4-(4-methylthiazol-5-yl)benzyl)pyrrolidine-2-carboxamide to get the compound **MS147N1** as a white solid in TFA salt form (14.6 mg, 65%). $^1$H NMR (600 MHz, Methanol-$d_4$) δ 9.07 (s, 1H), 8.43 (s,

1H), 8.11 (s, 1H), 7.85 – 7.80 (m, 2H), 7.48 – 7.44 (m, 3H), 7.44 – 7.40 (m, 2H), 7.16 – 7.10 (m, 2H), 6.40 – 6.35 (m, 2H), 4.84 (s, 2H), 4.59 (dd, $J$ = 8.4, 6.6 Hz, 1H), 4.54 – 4.47 (m, 3H), 4.40 (d, $J$ = 15.6 Hz, 1H), 4.02 – 3.92 (m, 3H), 3.77 – 3.67 (m, 3H), 3.29 – 3.22 (m, 4H), 3.17 – 3.10 (m, 3H), 2.51 (s, 3H), 2.33 – 2.20 (m, 4H), 2.15 (s, 1H), 2.11 – 2.04 (m, 1H), 1.86 – 1.79 (m, 2H), 1.72 (p, $J$ = 7.2 Hz, 2H), 1.55 – 1.42 (m, 4H), 1.30 – 1.24 (m, 4H), 1.09 (s, 9H). $^{13}$C NMR (151 MHz, Methanol-$d_4$) δ 175.4, 174.3, 173.0, 171.1, 153.2, 151.7, 151.5, 151.4, 150.7, 147.6, 145.4, 142.0, 141.9, 138.8, 132.0, 130.0, 129.0, 128.1, 127.3, 124.4, 115.6, 114.0, 110.0, 107.0, 69.1, 59.5, 58.6, 57.9, 55.4, 52.8, 48.4, 48.2, 42.1, 38.9, 37.7, 37.3, 35.5, 34.8, 34.0, 29.7, 28.9, 28.6, 26.3, 25.7, 25.6, 25.3, 23.6, 14.6. HRMS *m/z* $[M + H]^+$ calcd for $C_{54}H_{71}N_{12}O_6S^+$ 1015.5335, found 1015.5345.

**Scheme S2. The synthesis of MS147N2**

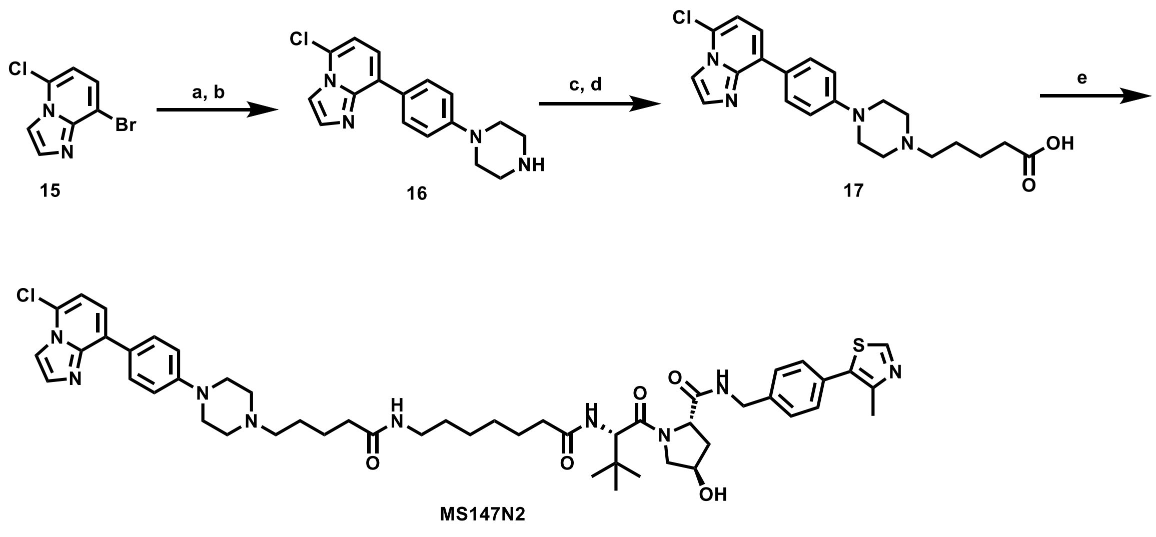


Reaction Condition: (a) 4-(4-*tert* Butoxycarbonylpiperazinyl)phenylboronic acid pinacol ester, Pd(dppf)$Cl_2$, $K_2CO_3$, dioxane, $H_2O$, 130 °C; (b) TFA, DCM, rt; (c) methyl 5-bromovalerate, $K_2CO_3$, DIEA, DMF, rt; (d) LiOH, MeOH, $H_2O$, THF; (e) EDCI, HOAt, NMM, DMSO, rt.

**5-Chloro-8-(4-(piperazin-1-yl)phenyl)imidazo[1,2-*a*]pyridine (16)**

**16** was synthesized following the same procedure as **12**. MS (ESI) *m/z* = 313.2 $[M + H]^+$.

**5-(4-(4-(5-chloroimidazo[1,2-*a*]pyridin-8-yl)phenyl)piperazin-1-yl)pentanoic acid (17)**

**17** was synthesized following the same procedure as **14**. $^1H$ NMR (600 MHz, Methanol-$d_4$) δ 8.47 (s, 1H), 8.21 (s, 1H), 8.03 – 7.97 (m, 2H), 7.58 – 7.53 (m, 2H), 7.25 – 7.20 (m, 2H), 4.13 – 3.95 (m, 2H), 3.81 – 3.65 (m, 2H), 3.30 – 3.24 (m, 3H), 3.23 – 3.11 (m, 2H), 2.46 – 2.42 (m, 2H), 1.92 – 1.83 (m, 2H), 1.78 – 1.69 (m, 2H). MS (ESI) *m/z* = 413.4 $[M + H]^+$.

**(2*S*,4*R*)-1-((*S*)-2-(7-(5-(4-(4-(5-chloroimidazo[1,2-*a*]pyridin-8-yl)phenyl)piperazin-1-yl)pentanamido)heptanamido)-3,3-dimethylbutanoyl)-4-hydroxy-*N*-(4-(4-methylthiazol-5-yl)benzyl)pyrrolidine-2-carboxamide (MS147N2)**

To a solution of compound **17** (8.3 mg, 0.02 mmol) in DMSO (1 mL), was added (2*S*,4*R*)-1-((*S*)-2-(7-aminoheptanamido)-3,3-dimethylbutanoyl)-4-hydroxy-*N*-(4-(4-methylthiazol-5-yl)benzyl)pyrrolidine-2-carboxamide (14.3 mg, 0.02 mmol, 1.0 equiv), EDCI (5.8 mg, 0.03 mmol, 1.5 equiv), HOAt (4.1 mg, 0.03 mmol, 1.5 equiv), and NMM (6.1 mg, 0.06 mmol, 3.0 equiv). The resulting mixture was stirred at room temperature overnight and then purified by preparative HPLC to get the compound **MS147N2** as a white solid in TFA salt form (5.6 mg, 24%).$^1H$ NMR (600 MHz, Methanol-$d_4$) δ 9.05 (s, 1H), 8.48 (d, *J* = 2.1 Hz, 1H), 8.23 (d, *J* = 2.2 Hz, 1H), 8.04 – 8.02 (m, 2H), 7.57 – 7.53

(m, 2H), 7.51 – 7.48 (m, 2H), 7.46 – 7.43 (m, 2H), 7.24 – 7.21 (m, 2H), 4.65 (s, 1H), 4.62 – 4.53 (m, 2H), 4.52 – 4.50 (m, 1H), 4.39 (d, *J* = 15.5 Hz, 1H), 4.06 – 4.01 (m, 2H), 3.92 (d, *J* = 11.1 Hz, 1H), 3.82 (dd, *J* = 10.9, 3.9 Hz, 1H), 3.76 – 3.69 (m, 2H), 3.32 – 3.24 (m, 5H), 3.24 – 3.15 (m, 4H), 2.36 – 2.21 (m, 6H), 2.13 – 2.07 (m, 1H), 1.88 – 1.80 (m, 2H), 1.76 – 1.69 (m, 2H), 1.68 – 1.60 (m, 2H), 1.56 – 1.50 (m, 2H), 1.42 – 1.33 (m, 5H), 1.05 (s, 9H). MS (ESI) *m/z* = 952.6 $[M + H]^+$.

### 1H NMR and 13C NMR spectra of key compounds

#### $^{1}H$ NMR and $^{13}C$ NMR spectra of MS147 in Methanol-$d_4$

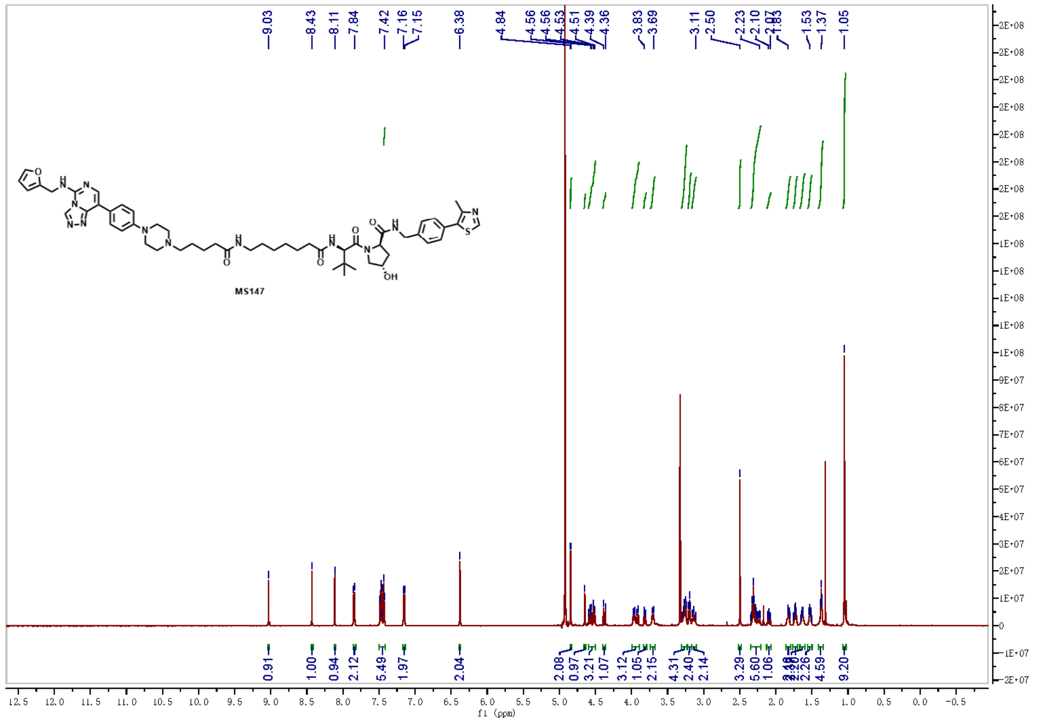

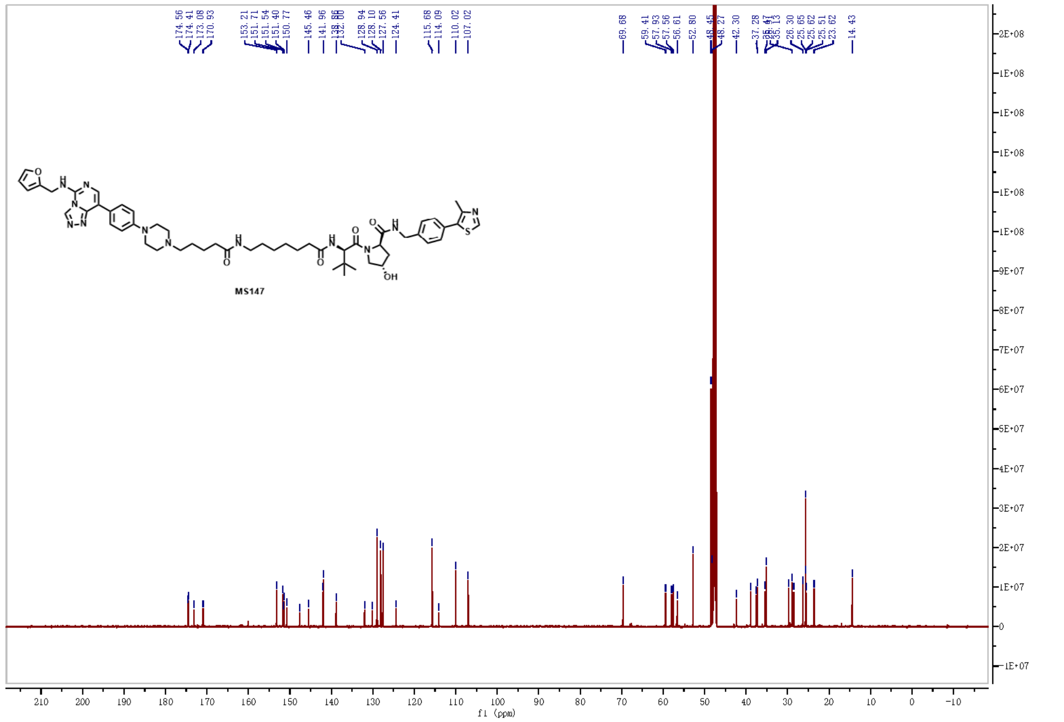

### $^{1}H$ NMR and $^{13}C$ NMR spectra of MS147N1 in Methanol-$d_4$

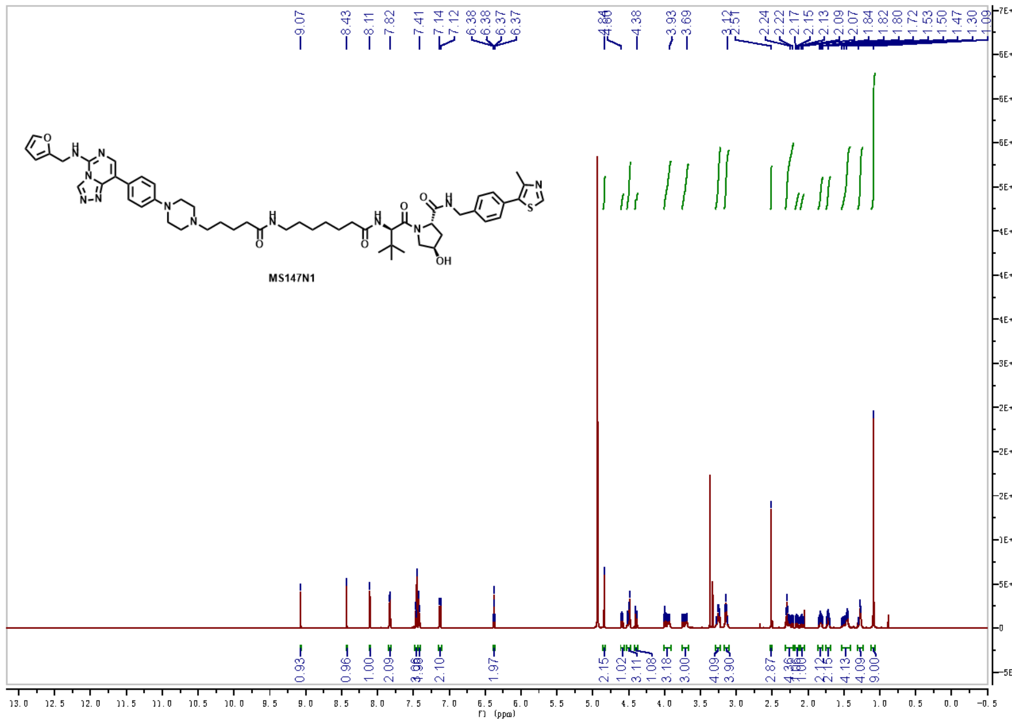

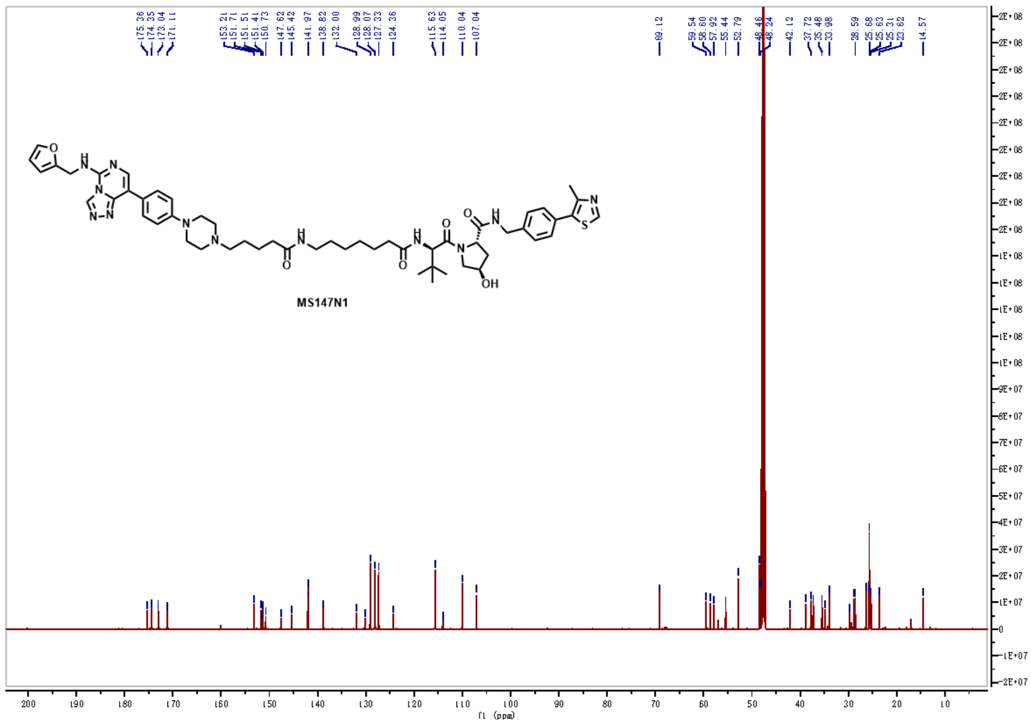

**[1]H NMR spectra of MS147N2 in Methanol-$d_4$**

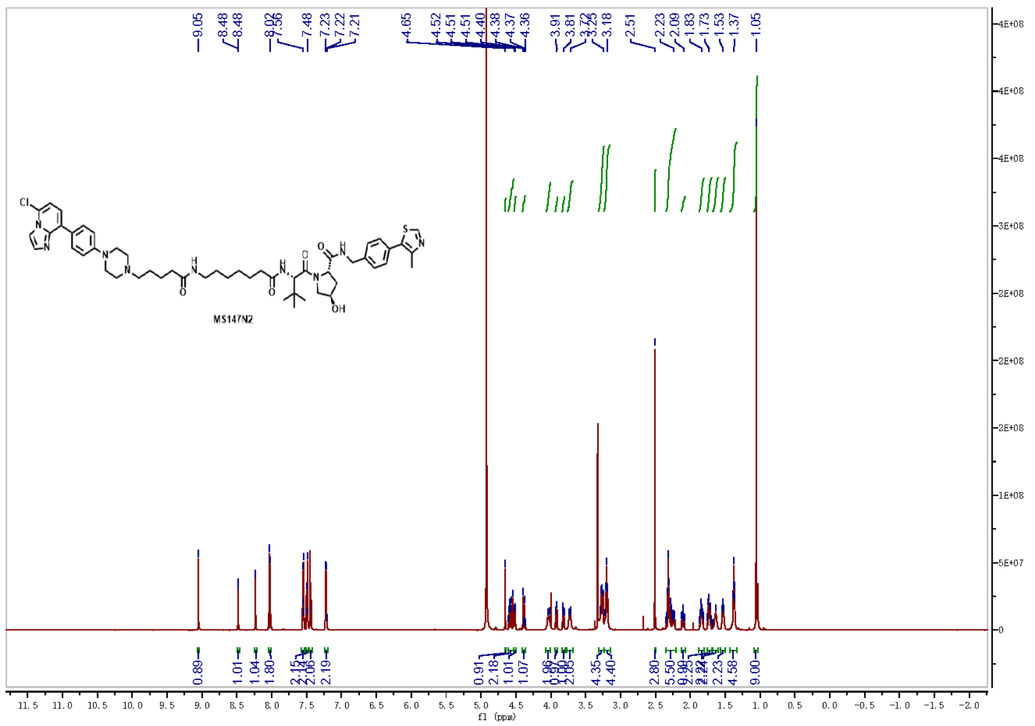
